# Loss of vitamin C biosynthesis protects from a parasitic infection

**DOI:** 10.1101/2025.07.22.666193

**Authors:** Gongwen Chen, Ji Hyung Jun, Tobias Wijshake, Yunyang Li, Minwei Yuan, Joseph Rose, Shan Li, Sarah Cobb, Willow Serpa, Yafeng Li, Li Li, Weina Chen, James J. Collins, Jipeng Wang, Michalis Agathocleous

## Abstract

The ability to synthesize essential molecules is sometimes lost in evolution. A classic example is ascorbate (Vitamin C), which is synthesized in most animals by L-Gulonolactone Oxidase (GULO), an enzyme lost multiple independent times in animal evolution. This event is thought to be evolutionarily neutral, however, *GULO-*deficient animals including humans need to obtain ascorbate from their diet and are susceptible to ascorbate deficiency and scurvy. We therefore hypothesized that this disadvantage of GULO loss is offset by physiological benefits. Here we show that ascorbate deficiency protects mice from schistosomiasis, a debilitating parasitic disease which afflicts 250 million people. *Schistosoma mansoni* worms required host ascorbate to produce eggs *in vivo.* Consequently, ascorbate-deficient mice were protected from schistosomiasis pathologies and transmission. Intermittent ascorbate deficiency protected *Gulo*-deficient mice from both scurvy and schistosomiasis mortality. The effects of ascorbate on schistosome reproduction were mediated by ascorbate-dependent histone demethylation which promoted vitellocyte development in female schistosomes. We propose that vitamin deficiencies are not always detrimental but can protect animals from pathogens which need to obtain vitamins from their host.

## INTRODUCTION

Ascorbate is a small molecule synthesized in most animals by an evolutionarily ancient metabolic pathway terminating in the enzyme L-Gulonolactone Oxidase (GULO)^1^. GULO, whose only known function is ascorbate synthesis, was lost repeatedly and independently in the evolution of many species, including haplorrhine primates, cavy rodents, fruit bats, teleost fishes, and passeriform birds^1^ (**Fig. 1a**). This loss transformed ascorbate into a vitamin, whose consumption is essential for health and survival in GULO-deficient species, including humans. It is assumed that loss of ascorbate synthesis incurs no fitness costs because it can be fully replaced by dietary intake, and *GULO* loss is a paradigmatic evolutionarily neutral gene loss^2^. However, unlike ascorbate-synthesizing organisms such as mice which maintain uniformly high systemic ascorbate levels, *GULO* loss allows for large variations in plasma ascorbate levels^3,4^. Low ascorbate levels in humans are common and are associated with increased mortality from noncommunicable disease^4,5^. Ascorbate is an antioxidant^6^ and stimulates the activity of many oxygenases^7^. Ascorbate deficiency impairs hematopoiesis^8^, germline development^9^, bone development^10^, norepinephrine synthesis^11^, collagen production, vascular integrity^12^, and other cellular processes^13^. Persistent deficiency causes scurvy, a fatal disease^14,15^. Why would some species lose the capacity for ascorbate synthesis if ascorbate has many important functions, and its deficiency causes disease? We hypothesized that ascorbate deficiency must confer some robust benefit to the organism in particular physiological settings that may outweigh its harms. Because GULO evolved in the earliest eukaryotes (**Fig. 1a**), any eukaryote which does not have it must have lost it during evolution and therefore may require exogenous ascorbate. This also applies to pathogens, some of which contain GULO or related genes and can synthesize their own ascorbate, and some of which cannot (**Fig. 1b**).

**Figure 1.**
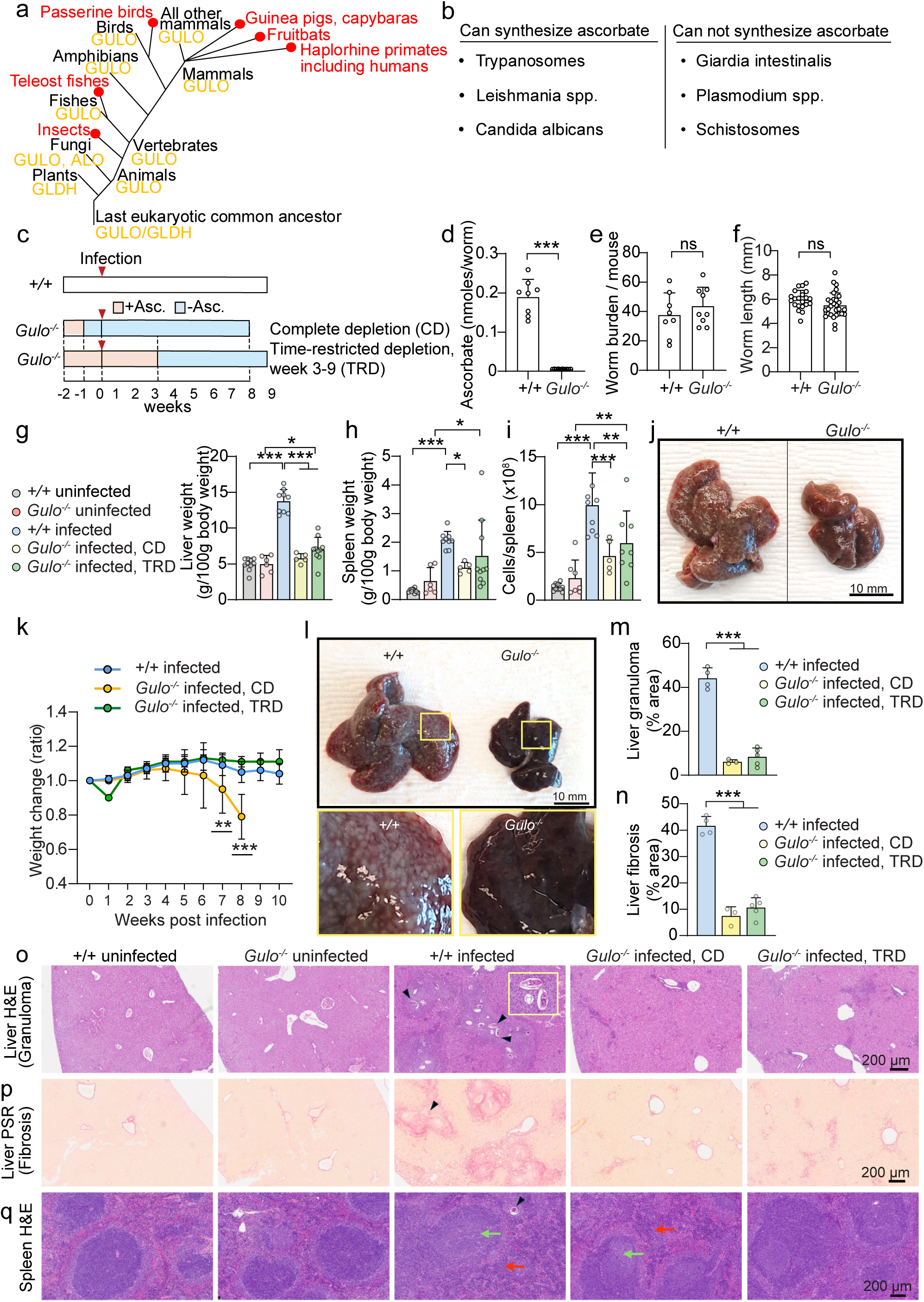
Ascorbate deficiency prevents schistosomiasis pathology. **a.** Ascorbate synthesis through L-Gulonolactone oxidase (GULO) or its paralogues GLDH/ALO evolved in the last eukaryotic common ancestor and was lost multiple independent times during animal evolution. **b.** Examples of human pathogens which can or cannot synthesize their own ascorbate (or ascorbate analogues such as erythroascorbate). **c.** *S. mansoni* infection of *Gulo^-/-^* mice. *Gulo^-/-^* mice were placed on complete ascorbate depletion (CD) starting from 1 week before infection or on time-restricted ascorbate depletion (TRD) starting from 3 weeks after infection. Wild type control mice synthesize their own ascorbate and were not provided with exogenous ascorbate. **d.** Ascorbate levels in worms isolated from infected wild type and *Gulo^-/-^* mice (n = 8 wild type and 10 *Gulo^-/-^* mice, 2 worms/mouse). **e-f**. Worm number (n = 8-9 mice/treatment) and length (n = 21-29 worms/treatment) isolated from wild type or *Gulo^-/-^* infected mice. **g-i**. Liver and spleen weights and spleen cellularity at 8 weeks or 9 weeks post-infection (n = 5-10 mice/treatment) **j.** Granulomas at 7 weeks post-infection in the liver of infected wild type but not *Gulo^-/-^* mice with complete ascorbate depletion. **k.** Effect of complete (CD) or time-restricted (TRD) ascorbate depletion on body weight after infection. N = 5-7 mice/treatment. **l.** Granulomas at 9 weeks post-infection in the liver of infected wild type but not *Gulo^-/-^*mice with time-restricted ascorbate depletion. Insets show a magnification of the boxed area. **m.** Area of H&E-stained liver sections occupied by granulomas. n = 3-5 mice/treatment. **n.** Area of Picrosirius Red (PSR)-stained liver sections which shows fibrosis. n = 3-5 mice/treatment. **o.** Representative images of H&E-stained liver sections. Infected wild-type but not *Gulo^-/-^*mice show severe granuloma formation associated with heavy egg lodging (black arrowhead). Inset shows a magnified region with egg deposition. **p.** Representative images of PSR-stained liver sections. Infected wild-type but not *Gulo^-/-^* mice show severe hepatic fibrosis marked by collagen deposition. **q.** Representative images of H&E-stained spleen sections. Infected wild-type mice show proliferation of white pulp with follicular hyperplasia/germinal centers (green arrow) and expanded red pulp (red arrow) with associated extramedullary hematopoiesis (EMH). These features are attenuated in infected *Gulo^-/-^* mice. All graphs show means ± sd. Statistical significance was assessed with Welch’s t-test (d), unpaired t-test (e-f), and one-way ANOVA (g-n). *P < 0.05, **P < 0.01, ***P < 0.001.

Schistosome parasitic flatworms infect 250 million people every year and cause significant mortality and morbidity in the world’s poorest regions^16,17^. Schistosomes live in the circulation for decades and lay hundreds or thousands of fertilized eggs every day^18,19^. This massive fecundity causes schistosomiasis pathology and transmission^18,19^. Eggs lodge in organs, causing inflammation and the chronic disabling symptoms of schistosomiasis. Eggs trigger granuloma formation, facilitating egg shedding from the body^20^ and parasite transmission to the freshwater snails which serve as intermediate hosts^18^. Praziquantel, the only available treatment, is partially effective, is ineffective against immature worms^21^, and its intensive use may cause resistance^22^. It is essential to understand the molecular mechanisms of schistosome reproduction in order to prevent pathology and transmission.

Schistosome reproduction is governed by a coordinated developmental program in which contact with a male in the host stimulates female maturation and production of oocytes and vitellocytes^23^. Vitellocytes provide nutrients to fertilized eggs and express proteins which are cross-linked to form the eggshell^24^. Eighty years ago it was reported that schistosome eggshell formation is disrupted in infected guinea pigs with scurvy but whether this confers benefits to the host has been unclear^25^. Addition of ascorbate to culture media allows production of viable eggs from *Schistosoma mansoni* and *Schistosoma japonicum*^26,27^. Therefore, we hypothesized that vitamin C deficiency protects animals from schistosomiasis by depriving schistosomes of vitamin C, preventing parasite reproduction, host pathology, and disease transmission.

## RESULTS

### Ascorbate deficiency prevents schistosomiasis liver pathology

To test if vitamin C deficiency protects from schistosomiasis, we infected wild-type mice, which can synthesize ascorbate endogenously, or ascorbate-deficient *Gulo^-/-^* mice with the human pathogen *S. mansoni*. *Gulo^-/-^* mice were fed an ascorbate-deficient diet starting from 1 week before infection to induce ascorbate deficiency throughout the experiment (**Fig. 1c**). Worms isolated from wild type mice contained ascorbate, but worms isolated from ascorbate-deficient *Gulo^-/-^* mice did not (**Fig. 1d**). Therefore, schistosomes obtain ascorbate from their host. The number and size of worms was not significantly different between wild type and *Gulo^-/-^*mice suggesting ascorbate was not required for schistosome survival or growth *in vivo* (**Fig. 1e, f**). Infected wild type mice had hepatomegaly and splenomegaly typical of schistosomiasis^28^ but infected *Gulo^-/-^* mice had no hepatomegaly and their splenomegaly was attenuated (**Fig. 1g-i**). Extensive granulomas were present on the surface of infected wild type but not *Gulo^-/-^*livers (**Fig. 1j**). In this depletion schedule, ascorbate-deficient *Gulo^-/-^* mice were weak and had weight loss consistent with development of scurvy (**Fig. 1k**). To test if ascorbate deficiency can block schistosomiasis pathology without causing scurvy, in parallel we provided *S. mansoni* infected *Gulo^-/-^*mice with 1% ascorbate in the diet – which normalizes plasma ascorbate levels^8^ - before and for up to 3 weeks after infection and then transferred to an ascorbate deficient diet until analysis at 8-9 weeks after infection (**Fig. 1c**). This time-restricted depletion schedule removed ascorbate specifically during the egg-laying period whose onset is 4-5 weeks after infection^19^. *Gulo^-/-^* mice in this treatment group did not show signs of scurvy, including weight loss, lethargy, or weakness (**Fig. 1k**). Similar to the complete depletion group, time-restricted ascorbate depletion prevented hepatomegaly and reduced splenomegaly (**Fig. 1g-i)**. *Gulo^-/-^* mice on a time-restricted ascorbate depletion had no viable eggs deposited in their livers and no or minimal granuloma formation and liver fibrosis, in contrast to wild type mice which had abundant egg deposition, granuloma formation, and liver fibrosis (**Fig. 1l-p**). Histological analysis of the spleen showed that infection in wild type mice induced white pulp expansion with follicular hyperplasia and red pulp expansion with extramedullary hematopoiesis (EMH), effects that were reduced in ascorbate-depleted *Gulo^-/-^*mice (**Fig. 1q**).

Provision of ascorbate to *Gulo^-/-^* mice throughout the infection period in the chow or water restored hepatomegaly (**Supplementary Fig. 1a**) consistent with ascorbate deficiency being responsible for the phenotype of infected *Gulo^-/-^* mice. Therefore, ascorbate depletion can protect *S. mansoni*-infected mice from egg deposition, granuloma formation, inflammation, liver fibrosis, hepatomegaly and splenomegaly in the absence of scurvy.

### Ascorbate deficiency prevents hematopoietic and immune aberrations caused by schistosomiasis

*S. mansoni* infection causes hematopoietic and immune aberrations, including eosinophila^19,28^. Complete or time-restricted ascorbate depletion blunted the infection-induced increase in blood, bone marrow, spleen, and liver eosinophils (**Fig. 2a-f**) and eosinophil precursors (**Fig. 2g-i**). Ascorbate depletion prevented increased blood monocytes in infected mice (**Fig. 2j**). Consistent with our prior results^28^, infection impaired the development of the erythroid and platelet lineages, as suggested by a decreased frequency of megakaryocyte-erythroid progenitors (MEPs) and an increased frequency of platelet/erythroid-biased hematopoietic progenitors (HPC-2)^29,30^. These effects were rescued by ascorbate depletion (**Fig. 2k-l**). Schistosomiasis severely disrupted bone marrow erythropoiesis^28^, and this was also rescued by ascorbate depletion (**Fig. 2m-o**).

**Figure 2.**
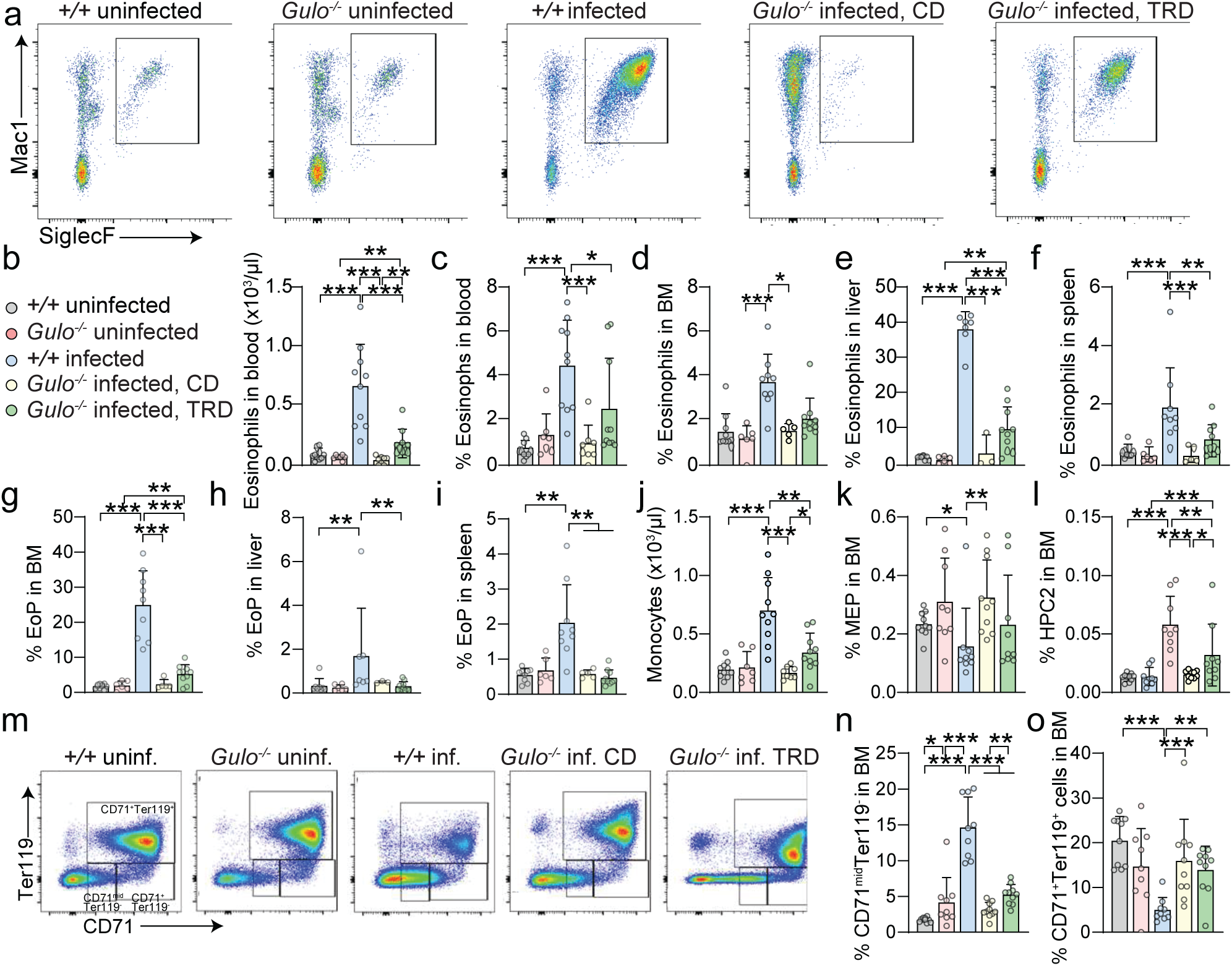
Ascorbate deficiency prevents hematopoietic aberrations in *S. mansoni* infected mice. **a**. Representative flow plots of liver eosinophils from uninfected wild type and *Gulo^-/-^*mice, or infected wild type and *Gulo^-/-^* mice on a time-restricted (TRD, weeks 3-9) or complete (CD) ascorbate depletion. Plots are gated out of CD45^+^CD115^-^LY6c^+/med^ cells. **b-c**. Number and frequency of eosinophils in the blood. n = 7-10 mice/treatment. **d-f**. The frequency of eosinophils in bone marrow (BM), liver and spleen. n = 3-10 mice/treatment. **g-i.** The frequency of SiglecF^+^Mac1^-^ eosinophil precursors (EoP) cells in bone marrow (BM), liver and spleen. n = 5-10 mice/treatment. **j**. The number of monocytes in the blood. n = 5-10 mice/treatment. **k**, **l**. The frequency of MEP and HPC-2 in bone marrow. n = 7-10 mice/treatment. **m**. Representative flow plots of erythrocyte progenitors quantified in n and o, gated out of Mac1^-^B220^-^ CD3^-^ cells. **n**, **o**. The frequency of CD71^mid^Ter119^-^ and CD71^+^Ter119^+^ erythroid progenitor cells. n = 9-10 mice/treatment. All graphs show means ± sd. Statistical significance was assessed with one-way ANOVA of log-transformed values (b, c, g, l), Kruskal-Wallis test (d, h, k), one-way Brown-Forsythe ANOVA (e, i, j), one-way ANOVA (f, o) and one-way Brown-Forsythe ANOVA of log-transformed values (n). *P < 0.05, **P < 0.01, ***P < 0.001.

Granulomas form around schistosome eggs a few days after deposition and continue to grow for several weeks^20,31^. To test if granuloma maintenance is affected by ascorbate, we supplemented *Gulo^-/-^* mice with ascorbate in the drinking water for one week during the egg laying period of the infection, followed by a 2-week ascorbate depletion and analysis (**Supplementary Fig. 1b**). In this setting, many granulomas at analysis were formed around eggs laid during the supplementation period. Ascorbate supplementation restored the effects of schistosomiasis on hepatosplenomegaly, granuloma formation, and hematopoiesis (**Supplementary Fig. 1c-n**). This suggests that ascorbate is not required for the maintenance of granulomas after they have formed. The number of schistosome eggs in the liver was reduced (**Supplementary Fig. 1o**) and eosinophilia was blunted (**Supplementary Fig. 1p-u**) after a 2-week ascorbate depletion, consistent with a reduction in ongoing egg deposition during the infection.

### Ascorbate deficiency protects from mortality and disease transmission

For ascorbate depletion to be protective against schistosomiasis, benefits to the host must outweigh the adverse effects of scurvy. Development of scurvy requires ascorbate depletion for several months in humans^32,33^, and for around 6 weeks in mice^12^ but schistosomes lay eggs daily. We hypothesized that the different timescales of the harms compared to the benefits of ascorbate deficiency might provide a survival advantage during infection. To assess this, we tested if intermittent ascorbate depletion for 3 weeks followed by repletion for 1 week may benefit the host by decreasing the intensity of egg laying and pathology while preventing scurvy. Infected *Gulo^-/-^* mice with *S. mansoni* were placed on this long-term intermittent ascorbate depletion schedule (**Fig. 3a**). 10/16 infected wild type mice died after infection, but only 1/19 of intermittently depleted *Gulo^-/-^*mice died (**Fig. 3a**). Serial bleeds revealed a reduction in eosinophilia and monocytosis in infected *Gulo^-/-^* as compared to wild type mice suggesting reduced inflammation (**Fig. 3b-c**). Infected wild type mice had thrombocytopenia which was rescued in intermittently depleted *Gulo^-/-^* mice (**Fig. 3d**). Therefore, intermittent ascorbate depletion provides a robust survival advantage to *Gulo^-/-^*mice infected with *S. mansoni*.

**Figure 3.**
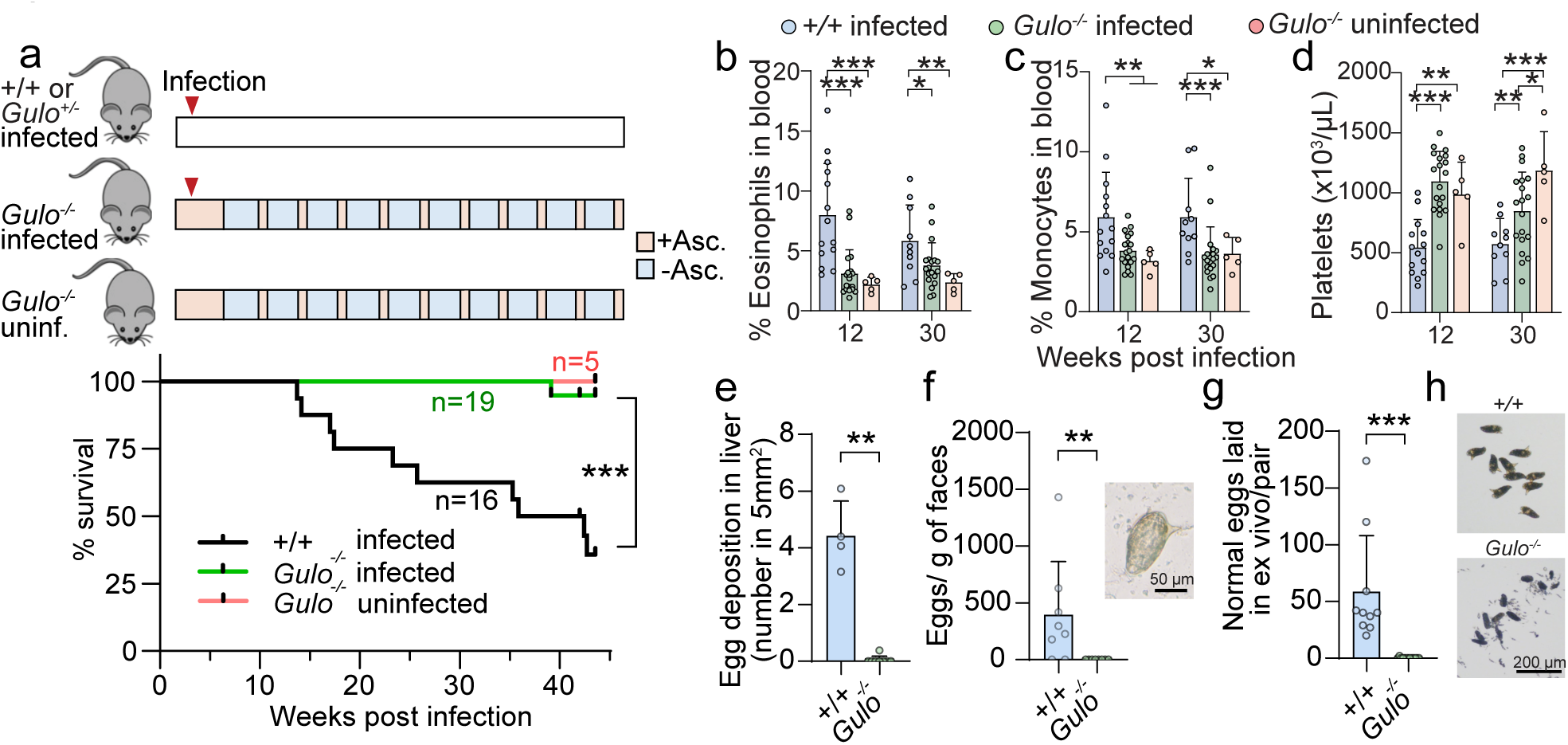
Ascorbate deficiency blocks schistosomiasis mortality, worm reproduction, and transmission. **a**. Schedule of *S. mansoni* infection with long-term alternating ascorbate depletion/repletion and Kaplan-Meier survival curve of wild type or *Gulo^-/-^* infected mice. *Gulo^-/-^*mice were supplied with ascorbate until 2.5 weeks after infection and then treated on a 3 week-depletion/1 week-repletion cycle. n=16-19 mice/treatment. 5 of 5 uninfected *Gulo^-/-^* control mice survived throughout the experiment. **b-d**. The frequency of eosinophils and monocytes in blood and platelet counts of infected wild type or *Gulo^-/-^* mice on the alternating depletion/repletion schedule. n = 5-19 mice/treatment. **e.** Egg deposition in liver of wild type or *Gulo^-/-^* mice that were either on a time-restricted (TRD, weeks 3-9) or a complete (CD) ascorbate depletion, analyzed at 8-9 weeks after infection. Egg number was counted from H&E-stained liver sections. n = 4 mice/treatment. **f.** Fecal egg load of infected wild type and *Gulo^-/-^* TRD mice, analyzed 9 weeks after infection. n = 8 mice/genotype. The image shows a miracidium recovered from feces of wild type mice. **g.** Rate of normal egg production per worm pair from worms isolated from infected wild type or ascorbate-depleted *Gulo^-/-^* mice. Egg number was counted at days 1 and 2 after isolating worms from mice and culturing in BM169 media without ascorbate. **h.** The image shows the representative developed miracidium recovered from feces of wild type mice. All graphs show means ± sd. Statistical significance was assessed with Mantel-Cox log rank test (a), one-way ANOVA of log-transformed values (b, c), one-way ANOVA (d) and Mann-Whitney test (e-g). *P < 0.05, **P < 0.01, ***P < 0.001.

Ascorbate is necessary for viable schistosome egg production *in vitro*^26,27^. We tested if host ascorbate is necessary for viable egg production *in vivo.* Eggs were often seen in the center of liver granulomas in wild type mice but not in the liver of *Gulo^-/-^* mice depleted from ascorbate during the egg-laying period (**Fig. 3e**). *S. mansoni* egg shedding in the feces transmits schistosomiasis^20^. Feces of wild type mice contained many normal eggs but feces of ascorbate depleted *Gulo^-/-^* mice contained no eggs (**Fig. 3f**), suggesting that host ascorbate deficiency blocks disease transmission. Worms isolated from wild type mice produced viable eggs *ex vivo* for at least 2 days^26,27^ but worms isolated from *Gulo^-/-^* mice produced only malformed, dead eggs (**Fig. 3g-h**).

To test if a different schistosome species also requires host-derived ascorbate for egg production and pathology, we analyzed *Gulo^-/-^* mice infected with *S. japonicum*. Adult *S. japonicum* females isolated from ascorbate-depleted *Gulo^-/-^* mice produced malformed eggs which adhered to each other, in contrast to females isolated from ascorbate-supplemented *Gulo^-/-^* mice which produced eggs with a normal morphology (**Supplementary Fig. 2a**). Ascorbate-depleted *Gulo^-/-^* mice were protected from *S. japonicum-*induced hepatomegaly, liver granulomas and fibrosis as compared to ascorbate-supplemented *Gulo^-/-^* mice (**Supplementary Fig. 2b-d**). Therefore, ascorbate deficiency protects against egg production and pathology of infection from multiple schistosome species.

### Ascorbate is required for female schistosome vitellocyte development

Our results suggest that host-derived ascorbate is essential for schistosome egg production, disease development and transmission. Ascorbate depletion did not affect parasite survival or growth *in vivo* (**Fig. 1e, f**), suggesting ascorbate specifically regulates reproduction rather than general growth processes. To understand how ascorbate regulates egg production, we first assessed the effects of ascorbate on female schistosome reproductive organ development. Maturation of the ovaries and vitellaria of virgin female schistosomes is triggered by pairing with a male schistosome ^23^. No changes were observed in the gross structure of ovaries of paired mature female *S. mansoni* isolated from ascorbate depleted *Gulo^-/-^*as compared to wild type mice (**Fig. 4a**) suggesting ovary maturation does not require ascorbate. In contrast, vitellaria maturation was inhibited in female *S. mansoni* isolated from ascorbate depleted *Gulo^-/-^*as compared to wild type mice (**Fig. 4a**). Female *S. japonicum* vitellaria development and egg production but not ovarian development was impaired in paired infections of ascorbate depleted as compared to ascorbate-supplemented *Gulo^-/-^* mice (**Supplementary Fig. 3**). These results suggests that ascorbate is required for female schistosome vitellaria maturation in the infected host.

**Figure 4.**
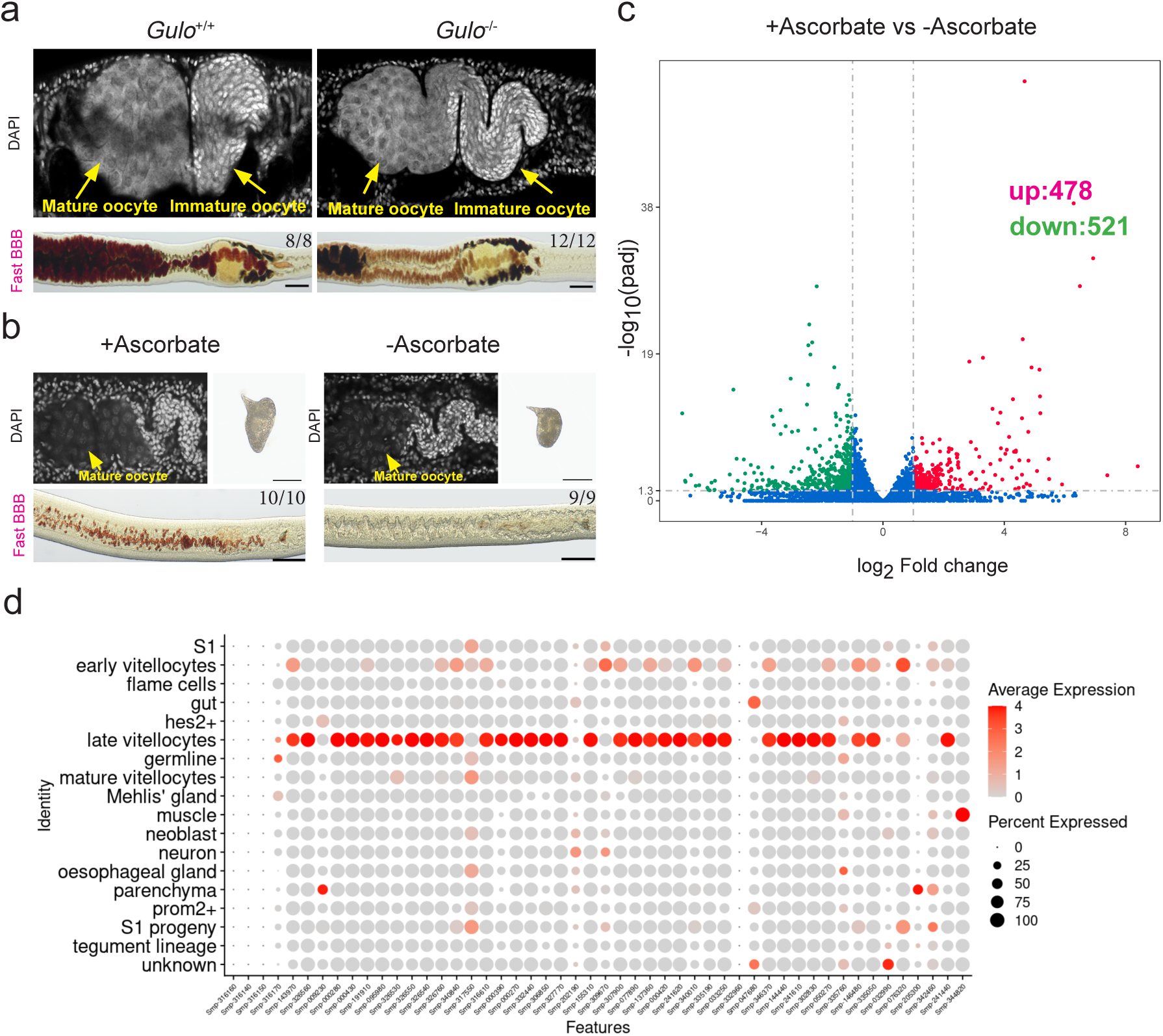
Host-derived ascorbate is essential for schistosome vitellaria development. **a.** DAPI and Fast BBB staining showing the vitellaria (top) and ovary (bottom) of adult female parasites from wild type mice or *Gulo^-/-^* mice. **b.** Reproductive organs and eggs laid by virgin female parasites cultured with BATT in the presence or absence of ascorbate. Vitellaria were labeled by Fast BBB staining, and nuclei were stained with DAPI. **c.** Volcano plot of the differentially expressed genes in virgin females treated with or without ascorbate. Genes were considered differentially expressed at Log_2_Fold change>1 and *padj* <0.05. **d.** Dot plot displaying the cell types which express the 50 most upregulated genes between ascorbate-supplemented and deprived conditions at the single-cell level.

We previously showed that ascorbate is crucial for egg formation in cultured worms^26^. To assess the mechanisms by which ascorbate promotes egg laying, we set up an *in vitro* system in which we could control the timing of female sexual development using the male peptide pheromone β-alanyl-tryptamine (BATT). BATT can trigger female sexual maturation in the absence of males^23^. Virgin females isolated from unisexual mouse infections were stimulated with BATT and cultured with or without ascorbate. After 10 days in culture, females cultured with ascorbate laid normal eggs, but those cultured without ascorbate produced smaller eggs with malformed or missing structures typical of normal eggs, such as a smooth external shell, a prominent lateral spine, and a viable oocyte (**Fig. 4b**). Vitellaria from ascorbate-treated virgin females were strongly stained with Fast Blue BB, indicative of vitellaria maturation, but untreated females had reduced staining indicative of impaired maturation (**Fig. 4b**). We did not observe differences in ovary maturation between the two groups (**Fig. 4b**). Therefore, the effects of ascorbate on female sexual maturation in the host could be recapitulated in this controlled developmental *in vitro* system.

To explore the molecular changes induced by ascorbate, we performed RNA-seq analysis on ascorbate-treated or untreated females stimulated with BATT. Ascorbate treatment changed the expression of many genes (**Fig. 4c, Supplementary Table 1**). To pinpoint the tissues affected by ascorbate at the transcriptional level, we mapped the gene expression changes induced by ascorbate onto a single-cell gene expression atlas we previously generated^34^. Most of the top 50 transcripts upregulated by ascorbate were specifically expressed in cells of the late vitelline lineage (**Fig. 4d**). Genes whose expression was upregulated by ascorbate included several involved in eggshell synthesis (**Supplementary Fig. 4**). These data suggest that female schistosomes depend on host-derived ascorbate to support vitellaria development.

To determine processes influenced by ascorbate treatment in an alternative way, we performed metabolomics in ascorbate-treated or untreated female schistosomes. The top metabolite enriched in ascorbate-treated schistosomes was L-DOPA, a product of tyrosinase activity, which crosslinks vitellocyte proteins to form the schistosome eggshell (**Supplementary Fig. 5, Supplementary Table 2**), consistent with a role for ascorbate in vitellocyte development.

### Molecular mechanisms of ascorbate action in female schistosomes

We investigated the molecular basis for the ascorbate requirement in egg production. Ascorbate is a major antioxidant^6^ but other antioxidants including vitamin E or glutathione could not replace ascorbate in promoting vitellaria development and egg production *in vitro* suggesting ascorbate does not act as an antioxidant (**Fig. 5a**). The action of ascorbate on egg production was stereospecific (**Fig. 5a**)^26^. Transport of the stereoisomer D-ascorbate into schistosomes was significantly lower than transport of the physiological L-ascorbate form (**Fig. 5b**). This suggested that the basis for stereospecificity in egg-induction was transport rather than enzymatic action, consistent with work in mammalian systems suggesting stereospecific transport as the basis for stereospecific ascorbate activity^35^. Ascorbate promotes the activity of many iron-and-α-ketoglutarate-dependent dioxygenases (α-KGDDs) in enzymatic assays, cell culture, and animals^7^. We therefore searched for dioxygenases which may be ascorbate targets in *S. mansoni*. Ascorbate activates TET enzymes which oxidize methylated cytosine (5mC) to produce 5-hydroxymethylcytosine (5hmC) on DNA^8,36–38^. Analysis with a sensitive mass spectrometry method^8^ did not detect 5mC or 5hmC in the schistosome genome (**Supplementary Fig. 6a, b**). AlkB-family α-KGDDs can demethylate 6-methyladenosine (m6A) on mRNA. Schistosome mRNA contained m6A (**Supplementary Fig. 6c**), however ascorbate supplementation did not significantly change its levels (**Supplementary Fig. 6d**). In mammals, ascorbate is a co-factor for the activity of collagen prolyl-4-hydroxylase^39^. However, neither total hydroxyproline levels nor newly synthesized hydroxyproline (**Supplementary Fig. 6e-f**) increased in ascorbate-treated schistosomes. Carnitine synthesis also relies on α-KGDD enzymes, but carnitine levels were not affected by ascorbate treatment (**Supplementary Fig. 6g**). Knockdown of *S. mansoni* orthologs of enzymes for nucleic acid modification or proline or lysine hydroxylation in paired female *S. mansoni* did not impair egg production (**Supplementary Fig. 7**).

**Figure 5.**
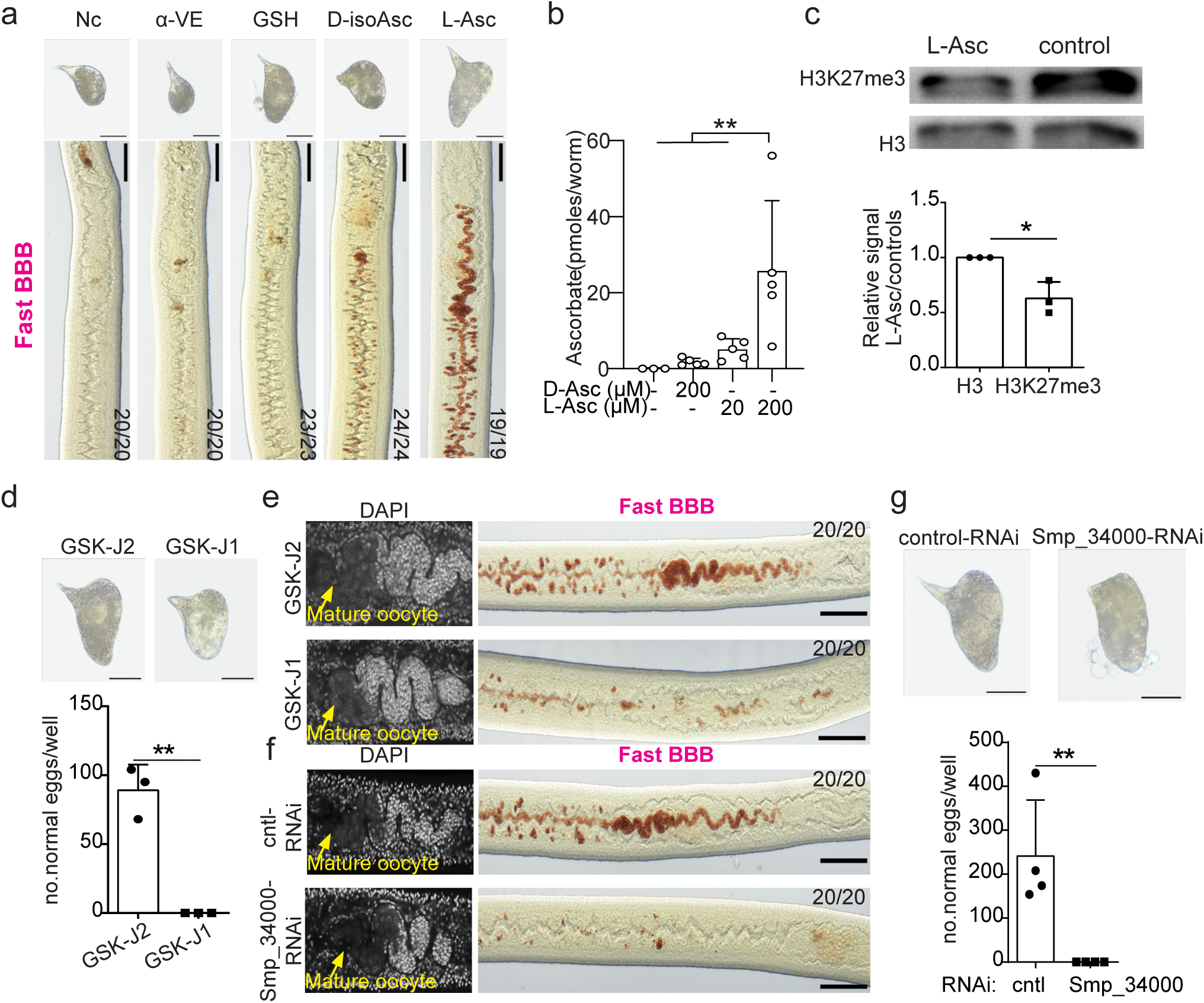
Ascorbate promotes schistosome reproduction via Kdm6. **a.** Eggs and vitellaria of virgin females treated with BATT in medium supplemented with different antioxidants. Vitellaria were labeled by Fast BBB staining. Nc= no treatment control; α-VE: vitamin E (alpha tocopherol); GSH=glutathione; D-isoAsc= D-isoascorbate; L-Asc=L-ascorbate. **b.** Ascorbate uptake in female worms. Paired worms were incubated in BM169 media containing L-ascorbate or D-ascorbate of the indicated concentrations for 4 days and ascorbate levels of single female worms were measured after separation. **c.** Evaluation of H3K27me3 levels in female parasites treated with or without ascorbate. Top, representative western blot image using the H3K27me3 antibody. Bottom, quantitation of H3K27me3 levels from Western blot images. **d.** Morphology and number of normal eggs per well laid by virgin females treated with BATT and either GSK-J2 or GSK-J1. **e.** DAPI and Fast BBB staining showing the ovary (left) and vitellaria (right) of virgin females treated with BATT and either GSK-J2 or GSK-J1. **f.** DAPI and Fast BBB staining showing the ovary (left) and vitellaria (right) of virgin females treated with BATT and either control or *Smp_034000* dsRNA. **g.** Morphology and number of normal eggs per well laid by virgin females treated with BATT and either control or *Smp_034000* dsRNA.

Jumonji C-domain histone demethylases are α-KGDDs whose activity is ascorbate-dependent in several systems, including purified enzyme assays, pluripotent stem cells *in vitro,* and animals^40^. Pharmacological inhibition of the Kdm6a/UTX H3K27me2/me3 demethylase homologue Smp_034000 inhibits schistosome egg production^41,42^. We thus tested if the effects of ascorbate in schistosome reproduction were mediated by histone demethylation. L-ascorbate treated BATT-stimulated female schistosomes had reduced levels of H3K27me3 as compared to untreated controls (**Fig. 5c**). The sequence of schistosome KDM6 JmjC domain has high homology to that of KDM6 in humans and other organisms (**Supplementary Fig. 8**) suggesting that inhibitors designed for human KDM6 could also be effective in schistosomes. Consistent with prior results^41,42^, pharmacological inhibition of H3K27me2/me3 demethylase activity with GSK-J4, an ester pro-drug of the KDM6 inhibitor GSK-J1^43^ inhibited normal egg production (**Supplementary Fig. 9a**). Ester conjugated pro-drugs may have spurious effects^44^. We therefore validated these results with treatment with GSK-J1, which can enter cells, albeit less efficiently than GSK-J4, and may be more selective for KDM6^45^. To avoid effects on male worms, we evaluated the action of GSK-J1 in BATT-stimulated virgin female *S. mansoni.* GSK-J1 did not cause gross morphological abnormalities in treated females but impaired vitellaria development at a dose of 6 µM (**Supplementary Fig. 9b**). GSK-J1-treated females produced fewer and abnormal eggs, as compared to females treated with the inactive control isomer GSK-J2 (**Fig. 5d**) and had impaired vitellaria but not ovary development (**Fig. 5e**). Similar to BATT-stimulated females, paired adult females treated with GSK-J1 also laid abnormal eggs compared to those treated with GSK-J2 (**Supplementary Fig. 9c**). To test if KDM6 genetically mediates the effects of ascorbate, we knocked down *Smp_034000* in females in the presence of ascorbate. Similar to the effects observed with GSK-J1 treatment, knockdown of *Smp_034000* inhibited ascorbate-induced vitellaria maturation and egg production, without impacting ovarian maturation (**Fig 5f, g, Supplementary Fig. 9d**). These results suggest that KDM6 inhibition phenocopies ascorbate deprivation in impairing egg production.

## DISCUSSION

Our work suggests that the enormous fecundity of the female schistosome and resulting morbidity, mortality, and disease transmission depend on ascorbate supply from the host. Ascorbate regulates mammalian stem cell function, cell differentiation, and reproduction via DNA or histone methylation^8,10,37,38^. Our finding that ascorbate regulates female *S. mansoni* reproduction via histone demethylation suggests that regulation of the epigenome by ascorbate is an evolutionarily generalizable mechanism for the control of cell differentiation and reproduction. Although our search did not reveal ascorbate targets aside from histone demethylation, we do not exclude the possibility that ascorbate may act via additional mechanisms to control schistosome egg production. More generally, our results support the idea that the abundance of a specific molecule in the host diet can affect the pathogenesis and transmission of parasitic disease by modulating parasite reproduction.

Ascorbate in the blood is depleted after several challenges including inflammation and infection^46–52^. Our results suggest that this depletion may be a mechanism to protect against pathogens which themselves require ascorbate. Ascorbate levels in the blood vary ten-fold in the U.S. and U.K.^4,5^. Ascorbate levels are low in schistosomiasis-endemic regions, suggesting ascorbate deficiency is physiologically relevant in at-risk populations^53^. In addition to schistosomes, many other pathogens have lost GULO during evolution and are thus likely unable to synthesize ascorbate^1^ (**Fig. 1b**). We predict that ascorbate deficiency protects against the pathogenesis of any infections in which the pathogen requires host ascorbate for its survival, development, or reproduction. Vitamin C supplementation should be avoided in such diseases.

Genes are often mutated or lost in evolution despite substantial costs to the organism because this protects from infection^54^. The widespread disease caused by vitamin C deficiency in diverse societies and contexts^14,15^ is hard to reconcile with the prevailing notion that over millions of years of evolution repeated independent *GULO* losses in many animal lineages have been evolutionarily neutral. Our work explains this paradox by demonstrating a physiological benefit of vitamin C deficiency. Schistosomes are generalist parasites that infect many species^55^ including most primates. Schistosomiasis or other helminth infections have been prevalent in untreated populations in endemic areas^56^, pre-modern societies^57^, and wild primates^58^, therefore it is possible these infections have exerted evolutionary pressures on the primate lineage. It remains possible that another pathogen drove *GULO* loss or that *GULO* loss resulted from neutral drift. In addition to vitamin C, the activity of several other vitamin synthesis or metabolism pathways has been lost or reduced in different species, and this has traditionally been ascribed to metabolic costs^59^. We propose the general idea that vitamin synthesis may be lost, or vitamin levels may be reduced as a protection mechanism against pathogens which obtain these vitamins from the host.

## METHODS

### *S. mansoni* parasites and mouse infections

For experiments performed at UT Southwestern, *S. mansoni* worm maintenance and infections were performed as described^23^. *S. mansoni* Naval Medical Research Institute (NMRI) strain was provided by the NIAID Schistosomiasis Resource Center. Infected *Biomphalaria glabrata* snails (M-line) and 6-7 week old female Swiss-Webster mice were used for in-house life cycle maintenance. Worm pairs were harvested from mice with mixed infections of male and female cercaria shed by several *B. glabrata* snails. For harvesting, adult worms were perfused from mice 7-9 weeks after infection using DMEM with 5% horse serum and heparin warmed at 37°C. Recovered worms were washed several times with perfusion solution and cultured in Basch Medium 169 containing 10% fetal bovine serum (BM169)^26^ supplemented with 1 x Antibiotic-Antimycotic (Sigma-Aldrich A5955, MO, USA). 5 worm pairs were cultured in 12-well plates containing 2 mL media per in 5% CO_2_ at 37°C. Media was changed every other day. To separate worm pairs, worms were suspended in BM169 with 0.25% tricaine and agitated for 5 min. Unpaired worms were processed further as indicated in the manuscript or washed two times in BM169 and returned to culture. For metabolomics, DNA and RNA analysis, and knockdown experiments, worms were cultured in ABC169 media^26^, with or without ascorbate. For experimental mouse infections, each mouse was infected with 200 *S. mansoni* cercariae released from infected *B. glabrata* snails by tail exposure. Mice were restrained, and their tail was dipped in water with cercariae for 30 min. Both male and female mice were used in all studies. Mice infected for experiments were on the C57BL/Ka background. Experimental wild type or *Gulo^-/-^* mice were bred in homozygous crosses on the C57BL/Ka-Thy-1.1 (CD45.2) or C57BL/Ka-Thy-1.2 (CD45.1) backgrounds, with the exception of long-term intermittent supplementation experiments to assess survival, for which *Gulo^-/-^* mice or their littermate *Gulo^+/-^*controls were bred from heterozygous (*Gulo^+/-^*) crosses. Ascorbate plasma concentrations of *Gulo^+/-^*mice are similar to those in wild type (*Gulo^+/+^*) mice^12^. *Gulo*^−/−^;CD45.1 mice were originally generated by crossing *Gulo*^−/−^ mice with C57BL/Ka-Thy-1.2 mice. *Gulo^-/-^* mice were maintained on 1% ascorbate-supplemented chow (Envigo) and switched to a standard mouse diet for ascorbate depletion before or after infection as noted in the results. In experiments with short-term intermittent ascorbate depletion (Supplementary Figure 1), mice were infected at 9-13 weeks of age. In experiments with time-restricted or complete ascorbate depletion, mice were infected at 14-21 weeks of age. In survival experiments with long-term alternating ascorbate depletion/repletion cycles, mice were infected at 23-32 weeks of age. Mice were analyzed 7-9 weeks after infection unless otherwise noted. In ascorbate supplementation periods, *Gulo^-/-^*mice were fed 1% ascorbate-supplemented chow unless otherwise noted. For *Gulo^-/-^*mice in the complete depletion (CD) treatment which developed scurvy, 20 mg/L ascorbate in the drinking water was provided in the final days of the experiment, a low supplementation dose lower than those needed to replenish plasma ascorbate^8^ or rescue schistosomiasis pathology. Mice were housed in the Animal Resource Center of University of Texas Southwestern Medical Center and all procedures were approved by the UT Southwestern Institutional Animal Care and Use Committee (IACUC) (approval APNs: 2017-102092, 2018-102427, 2020-102931).

For experiments performed at Fudan University, female BALB/c mice (6 weeks old, 16–24 g) were purchased from Beijing Vital River Laboratory Animal Technology Co., Ltd. (China) and used for *S. mansoni* maintenance. *Gulo^-/-^* mice on a C57BL/6JGpt background were generated by GemPharmatech (Jiangsu, China) using CRISPR/Cas9 technology. Three-week-old female *Gulo^-/-^* mice were provided with 50 mg/L L-ascorbic acid (Sigma-Aldrich A0278) in the drinking water for one week, with the solution changed twice per week. At four weeks of age, they were infected with cercariae (*Schistosoma mansoni*, 300 cercariae; *Schistosoma japonicum*, 40 cercariae). After *S. mansoni* infection, *Gulo^-/-^* mice remained on 50 mg/L L-Ascorbic acid drinking water for one week and then divided into an L-ascorbic acid treatment group, which continued to receive the same dose of ascorbate acid, and a control group which received water without ascorbate acid until analysis at seven weeks post-infection. For *S. japonicum* infection, *Gulo^-/-^* mice treated with or without 50 mg/L L-ascorbic acid were analyzed at six weeks post-infection. To obtain virgin female worms or adult paired *S. mansoni*, female BALB/c mice were infected with either female-only or mixed-sex *S. mansoni* cercariae. At seven weeks post-infection, mice were perfused through the hepatic portal vein using 0.9% sodium chloride solution (plus 200–350 U/mL heparin). Parasites were washed three times with BM169 medium before being transferred to the appropriate culture medium for in vitro cultivation. Experiments with animals at Fudan University were conducted in accordance with the guidelines for the Care and Use of Laboratory Animals of the Ministry of Science and Technology of the People’s Republic of China (2006398) and were approved by the Animal Care and Use Committee of Fudan University (Fudan IACUC 2021JS0078).

### Egg counts in the feces

Feces from each infected mouse were collected from the intestine after mouse euthanasia and feces weight was measured. After homogenization in 4% formaldehyde/phosphate buffered saline (PBS), the mixture was stored at room temperature, and eggs present in 100 μL were counted using a light microscope.

### Cell isolation and hematopoietic analysis

Hematopoietic analysis was performed as previously described^28^. Bone marrow cells were obtained by flushing femurs and tibia with a 25G needle in staining medium consisting of Ca^2+^/Mg^2+^-free Hank’s balanced salt solution (HBSS; Gibco), supplemented with 2% heat-inactivated bovine serum (Gibco). Spleens were mechanically dissociated by crushing between frosted plates and trituration in staining medium. Livers were enzymatically digested for 30 minutes at 37°C in 1.5 ml RPMI-1640 (Sigma), containing 250 μg/ml liberase (Roche) and 100 μg/ml DNase I (Roche). Cell suspensions were filtered through a 40 μm strainer. Cell number was assessed with a Vi-CELL cell viability analyzer (Beckman Coulter). Blood was collected by cardiac puncture using a 25G needle or by tail vein bleeds and mixed in a tube containing 5 μl 0.5M EDTA. Complete blood cell counts were determined using a hemavet HV950 (Drew Scientific). For flow cytometric analysis of blood, 40 μl was lysed in 1 ml of ammonium chloride buffer (ACK; 155mM NH_4_Cl; 10 mM KHCO_3_; 0.1 mM EDTA) for 10 minutes at 4°C. Cells were incubated with fluorescently conjugated antibodies for 90 minutes on ice when using CD34 antibody or for 30 minutes at 4^0^C. Cells were washed with staining media and resuspended in staining media containing 1 μg/ml DAPI or 1 μg/ml propidium iodide for live/dead discrimination. Cell populations were defined with the following surface markers: CD150^+^CD48^+^Lineage^-^Sca-1^+^Kit^+^ hematopoietic progenitor cells (HPC-2), CD34^-^CD16/32^-^Lineage^-^Sca-1^-^Kit^+^ megakaryocyte–erythroid progenitors (MEPs), Mac1^+^CD115^-^Ly6C^mid^/^+^SiglecF^+^ eosinophils, Mac1^-^SiglecF^+^ eosinophil precursors, Mac1^-^B220^-^CD3^-^ CD71^mid^Ter119^-^ immature erythroid progenitors, Mac1^-^B220^-^CD3^-^CD71^+^Ter119^-^ erythroid progenitors, and Mac1^-^B220^-^CD3^-^CD71^+^Ter119^+^ erythroid progenitors. The lineage cocktail for progenitor analysis consisted of CD2, CD3, CD5, CD8, Gr1, B220, Ter119. Analysis was performed using the LSRFortessa flow cytometer (BD Biosciences) or FACSCanto (BD Biosciences). Data were analyzed using FlowJo (Flowjo LLC) or FACSDiva (BD Biosciences).

### Schistosome drug treatments

Female schistosomes were cultured in BM169 medium supplemented with 50 µM BATT on a 6-well plate. Each well contained 6 mL of culture medium with either 10 pairs of adult worms or approximately 30 virgin females. Antioxidants were used at a final concentration of 0.2 mM for L-Ascorbic acid and D-isoascorbic acid (Sigma-Aldrich), 0.23 mM for α-Vitamin E, and 0.2 mM for L-Glutathione. GSK-J1 (6HY-15648, MedChemExpress, USA) or GSK-J2 (HY-15648A, MedChemExpress, USA) were used at a final concentration of 6 µM. Culture media were replaced every other day. Each treatment included at least three biological replicates. Samples were collected after 14 days of culture for paired worms or 12 days of culture for virgin females.

### RNA interference (RNAi)

dsRNA production and RNAi treatment were performed as previously described^60,61^. Oligo sequences used to generate dsRNA templates are listed in Supplementary Table 3. For RNAi experiments in Supplementary Fig. 7, worm pairs were treated with 50 µg/mL dsRNA in BM169 medium containing 200 μM ascorbic acid and 0.2% V/V bovine washed RBCs. The medium containing dsRNA was changed every two days for up to 20 days before analysis. Controls were treated with dsRNA transcribed from pJC53.2. To quantify egg production, eggs were removed from the culture media at each media change. Two days later, egg number and the number of worm pairs were counted to calculate daily egg production. For RNAi experiments in Fig. 5, approximately 30 virgin females were cultured in a 6-well plate in wells containing 6 mL of BM169 medium supplemented with 50 µM BATT. Worms were treated with 50 µg/mL dsRNA for target genes or control. The culture medium, BATT and dsRNA were changed every other day. Each treatment included at least three biological replicates. Parasites were collected after 12 days of culture.

### Western Blot

To collect female parasites, freshly perfused or cultured adult paired worms were separated by incubation in BM169 medium containing 0.25% anesthetic (tricaine) for 5 mins. Then the female worms were washed twice with PBS, and histones were extracted using the EpiQuik Total Histone Extraction Kit (OP-0006-100, Epigentek, USA) according to the manufacturer’s protocol. The extracted histones were subjected to SDS-PAGE. Proteins were transferred onto a PVDF membrane (Millipore) and incubated overnight at 4°C with primary antibodies (anti-Histone 3: PTM BIO, PTM-1002RM; anti-H3K27me3: PTM BIO, PTM-5002). After a 1-hour incubation with secondary antibodies (Anti-Mouse IgG: KPL, 5220-0341; Anti-Rabbit IgG: KPL, 5220-0336), protein bands were visualized using Fully Automated Chemiluminescence Imaging Analysis System (TANON, 5200). Western blot band intensities were analyzed using ImageJ, with Histone H3 serving as a reference for the relative quantification of histone methylation modifications of interest.

### Quantitative RT-PCR

Following dsRNA treatment, the female worms were separated from males with 0.25% tricaine in BM169. Each biological replicate consisted of 5 worms. RNA was extracted using TRIzol LS reagent (Life Technologies, 10296010) or Zymo Direct-zol kit (Genesee Scientific 11-331, CA, USA) and reverse transcribed with iScript (Bio-rad 1708891, CA, USA). Real time quantitative PCR was performed with an iTaq Universal SYBR Green Supermix (Bio-Rad, 1725122) and a CFX384 real-time system (Bio-Rad). The data were analyzed using Bio-Rad CFX Maestro software. Cytochrome C oxidase I (Smp_900000) was used as an endogenous control for normalization.

### Histological evaluation

Livers and spleens from control and *S. mansoni–*infected mice were collected and fixed in 10% formalin for at least 24 h. After fixation, the tissues were paraffin embedded, sectioned, and stained with H&E or PSR (Picrosirius red staining) by the UT Southwestern Histo Pathology Core. H&E sections of livers and spleens from uninfected control and infected mice were evaluated for pathology, hematopoiesis, erythropoiesis, and inflammation by a pathologist (WC). Images were acquired on a Hamamatsu Nanozoomer S60 slide scanner and egg length was measured with Image J. Fast Blue BB and DAPI staining was performed as previously described^26^ and imaged on a Olympus BX51 Microscope, Zeiss AxioZoom V16 Microscope or Nikon A1 Laser Scanning Confocal Microscope.

### RNA-seq

Virgin females were cultured in BM169 medium supplemented with 50 µM BATT with or without added ascorbate at a final concentration of 0.2 mM. Each well contained 6 mL medium and 30 females. Culture medium and ascorbate were replaced every other day. At day 10, the females were collected and stored at -80 ^0^C for RNA sequencing at Novogene (China) on the Illumina PE150 platform (Illumina, USA). Each group included 3 replicates. Quality control of the raw sequencing data was performed using the FASTQC program (http://www.bioinformatics.babraham.ac.uk/projects/fastqc/). Differential expression analysis was performed using DESeq2 and the differentially expressed genes (DEGs) were defined with adjusted padj < 0.05 and |log2FC|>1.

### Ascorbate measurement

For ascorbate measurements, 25 μl of a chilled solvent of 25:25:50 methanol:acetonitrile:water (v:v:v) containing 10 mM EDTA and an internal standard of 100 pmoles ^13^C-ascorbate per female worm was used for extraction. The samples were ground with a pestle, sonicated for 1 min, vortexed for 1 min and centrifuged at 20,000 × *g* for 15 min at 4 °C. The supernatant was diluted using 25 µL of water and analyzed using an AB Sciex 6500+ QTrap mass spectrometer coupled to a Shimadzu UHPLC system operating in negative mode to detect the transitions 175/115 (ascorbate) and 176/116 (^13^C-ascorbate) as previously described^8^.

### Metabolomics and mass spectrometry

Metabolites were extracted by adding 30 μl of chilled 40:40:20 methanol:acetonitrile:water (v:v:v) solvent to five female worms. The samples were ground with a pestle, sonicated for 1 min, vortexed for 1 min and centrifuged at 20,000 × *g* for 15 min at 4 °C. For metabolomics analysis, the supernatant was transferred to a new tube, centrifuged at 21,000 × *g* for 15 min at 4 °C and transferred to LC-MS autosampler vials. For proline hydroxylation analysis, the pellet was resuspended in 100 µL 6N HCl, sonicated for 15 min and incubated for 6 hrs at 100 °C. The solvent was removed in a vacuum concentrator (SpeedVac) and the residue was resuspended in 50 μl water. Samples were sonicated in 15 min and filtered with a Nanosep 3k column (PALL OD003C33, Fisher Scientific, MA, USA). Liquid chromatography was performed using a Millipore ZIC-pHILIC column (5 mm, 2.1 × 150 mm) with a binary solvent system of 10 mM ammonium acetate in water, pH 9.8 (solvent A) and acetonitrile (solvent B) with a constant flow rate of 0.25 mL/min. The column was equilibrated with 90% solvent B. The liquid chromatography gradient was: 0–15 min linear ramp from 90% B to 30% B; 15–18 min isocratic flow of 30% B; 18–19 min linear ramp from 30% B to 90% B; 19–27 column regeneration with isocratic flow of 90% B. Mass spectrometry was performed with a Thermo Scientific QExactive HF-X hybrid quadrupole orbitrap high-resolution mass spectrometer (HRMS) coupled to a Vanquish UHPLC as previously described^62^. Retention time information obtained by running chemical standards was used for metabolite identification. A pooled quality control (QC) sample was interspersed through the LC-MS run to monitor the quality and consistency of detection.

### Measurement of 5hmC, 5mC and 6mA by LC–MS/MS

To measure 5mdC and 5hmdC, genomic DNA was purified from female worms using Zymo Quick-DNA Miniprep kit (D3024, CA, USA) according to the manufacturer’s instructions. 1 μg of DNA was digested to nucleosides with DNA Degradase Plus (Zymo Research) with the following internal standards added to the mix: ^13^C-^15^N2-2-deoxycytidine (^13^C-^15^N2-2dC), D3-5-methyl-2-deoxycytidine (D3-5mdC), D3-5-hydroxymethyl-2-deoxycytidine (D3-5hmdC) (Toronto Research Chemicals). To measure 6mA, total RNA was purified using Zymo Direct-zol kit and mRNA was purified using Magnetic mRNA Isolation Kit (NEB, S1550S). mRNA was digested by 10 unit of nuclease P1 (NEB, M0660S) in 25 μl buffer for 2 h at 37 °C. 3 μl of 1M ammonium carbonate, 0.05 units of venom phosphodiesterase I (Sigma-Aldrich, P3243), and 5 units of CIP alkaline phosphatase (NEB, M0290S) were added to the reaction and incubated for 4 h at 37 °C. To calculate m6A, internal standards (Adenosine-^13^C5 and N6-Methyladenosine-^13^C) (Toronto Research Chemicals) were added in the reaction. Hydrolyzed DNA and RNA samples were filtered with a Nanosep 3k column to remove enzymes, and filtrates were analyzed by LC–MS/MS. For 5mdC and 5hmdC, liquid chromatography was carried out with a Scherzo SM-C18 (Imtakt) reverse phase C18 column, 3μm, 2.1× 100mm. Mobile phase A was 0.1% formic acid in water, and mobile phase B was 0.1% formic acid in acetonitrile. The gradient was: 0–2.5min 0% B; 2.5–24min 0–100% B; 24–25min 100% B; 25–27min 100–0% B; 27–20min 0% B. The flow rate was 0.5 ml/min. The flow was diverted to the waste for the first 1min. Mass spectrometry was performed with an AB Sciex QTRAP 6500+ or 7500 operating in MRM mode. Mass spectrometry parameters were optimized using pure standards. To measure 6mA, liquid chromatography was carried out with a Kinetex F5 reverse phase column, 2.6μm, 2.1× 150mm. Mobile phase A was 0.1% formic acid in water, and mobile phase B was 0.1% formic acid in acetonitrile. The gradient was: 0–3min 0% B; 3–24min 0–95% B; 24–26min 95% B; 26–26.2min 95–0% B; 26.2–30min 0% B. The flow rate was 0.25 ml/min. The flow was diverted to the waste for the first 0.5min. Mass spectrometry was performed with an AB Sciex QTRAP 7500 operating in MRM mode. Q1 and Q3 were set at unit resolution. Mass spectrometry parameters including entrance potential (EP), collision energy (CE), and collision exit potential (CXP) were optimized use commercially available pure standards. The following transitions were monitored in positive mode: 2dC 228.1/112.1; 5mdC 242.1/126.1; 5hmdC 258.1/142.1; ^13^C-^15^N2-2dC, 231.1/115.1; D3–5mdC 245.1/129.1; D3–5hmdC 261.1/145.1; A 268.1/136.1; m6A 282.1/150.1; ^13^C5-A 273.1/136.1; ^13^C1-m6A 283.1/151.1. Chromatogram peak areas were integrated using Multiquant (AB Sciex). The amount of each nucleoside in a sample was calculated by dividing the peak area of the endogenous nucleoside by the peak area of the internal standard.

### Mass spectrometry data analysis

For data acquired with the AB Sciex 6500+ or 7500, chromatogram peak areas were integrated using Multiquant (AB Sciex). For data acquired with the QExactive, chromatogram peak areas were integrated using Tracefinder (Thermo Scientific) using an in-house database. For metabolomics analysis, the peak area for each metabolite was normalized to the median peak area of all the metabolites in each sample. Data analysis of the processed raw values was performed with MetaboAnalyst or Graphpad Prism.

### Statistical analysis

Statistical analysis was performed with GraphPad Prism. To test if data significantly deviated from normality, a Shapiro-Wilk test was used 3 ≤ n < 20 or the D’Agostino & Pearson test when n ≥ 20. If the data did not significantly (p < 0.01 for at least one treatment) deviate from normality, we used a parametric test, otherwise data were log-transformed and tested for a significant deviation from normality. If the log-transformed data passed normality, a parametric test was used on the transformed data. If both the untransformed and log-transformed data did not pass the normality test, a non-parametric test was used on the untransformed data. To test if variability significantly differed among treatments, we used the F-test (for experiments with two treatments) or the Brown–Forsythe test (for experiments with more than two treatments). To assess the statistical significance of a difference between two treatments, we used a t-test for data that was normally distributed and had equal variability, or a t-test with Welch’s correction for data that was normally distributed and had unequal variability, or a Mann-Whitney test for data that was not normally distributed. To assess the statistical significance of a difference between more than two treatments, we used a one-way ANOVA for data which did not significantly deviate from normality and did not have significantly unequal variances, a Brown–Forsythe ANOVA for data which did not significantly deviate from normality and had significantly unequal variances, and a Kruskal–Wallis test for data which significantly deviated from normality.

## Acknowledgements

We thank Lu Zhao, George Wendt, and other members of the Collins lab for help with infections; the Moody Foundation Flow Cytometry Facility for flow cytometry; the UTSW Histo Pathology Core for sectioning and H&E staining; and the CRI Metabolomics core for mass spectrometry analysis. We thank S. Morrison and H. Zhu for comments on the manuscript.

## Author contributions

The group at Fudan University generated data in Figures 4, 5a, 5c-g, Sup. Figures 2-4, 8-9. The group at UT Southwestern generated data in Figures 1-3, 5b, Sup. Figures 1, 5-7. JHJ, JCC, JW, MA designed experiments; GC, JHJ, TW, YuL, MY, JR, SL, SC, WS, YaL, LL, WC, JW, MA performed experiments or contributed methodology; JCC, JW, MA acquired funding; JW, MA directed the project; MA wrote the manuscript; JHJ, JW edited the manuscript.

## Data and materials availability

Data are available in the main text or the supplementary materials.

---

**Supplementary Figure 1.**
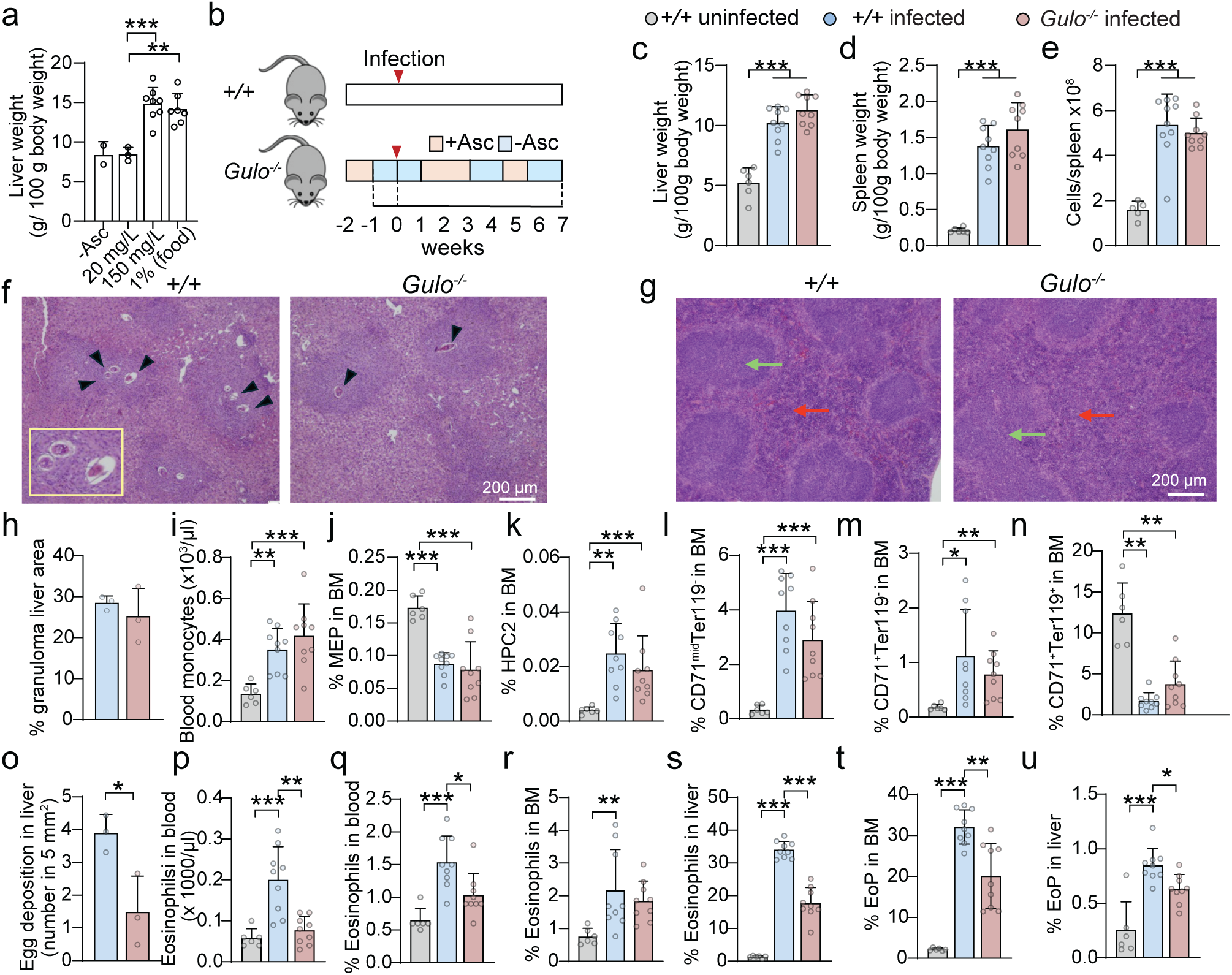
Effects of ascorbate supplementation on *S. mansoni-*infected *Gulo^-/-^*mice. **a.** Liver weight from *Gulo^-/-^* mice supplied with 20 mg/L or 150 mg/L ascorbate in the water or 1% ascorbate in the food at 7-8 weeks post-infection. n = 2∼8 mice/treatment. **b.** *S. mansoni* infection with short-term intermittent ascorbate supplementation during the egg laying period. Ascorbate was supplemented at 100-300 mg/L in the drinking water, changed every two days. **c-e.** Liver, spleen weight and spleen cell number at 7 weeks post-infection in wild type or *Gulo^-/-^* mice supplemented as in (b). n = 6-9 mice/treatment. **f.** Representative images of H&E-stained liver sections from infected wild type and *Gulo^-/-^*mice supplemented as in (b). Black arrowheads and magnified inset show heavier egg lodging in wild type as compared to *Gulo^-/-^* mice. **g.** Representative images of H&E-stained spleen sections from infected wild type and Gulo^-/-^ mice supplemented as in (b). Infected mice show proliferation of white pulp with follicular hyperplasia/germinal centers (green arrow) and significantly expanded red pulp (red arrow) with associated extramedullary hematopoiesis (EMH). **h.** Area of H&E-stained liver sections occupied by granulomas. n = 3 mice/treatment. **i-n.** The frequency of monocytes in the blood, MEP, HPC-2, and erythroid progenitors in the bone marrow. n = 6-9 mice/treatment **o.** Egg deposition in the liver. Egg number was counted from H&E-stained liver sections. n = 3 mice/treatment. **p-s.** Eosinophil blood cell count, blood, bone marrow, and liver frequency. n = 6-9 mice/treatment **t-u.** Eosinophil precursor (EoP) bone marrow and liver frequency. n = 6-9 mice/treatment All graphs show means ± sd. Statistical significance was assessed with one-way ANOVA Brown-Forsythe (a), one-way ANOVA (c-e, i-l, r, u), unpaired t-test (h, o), welch’s t-test (m, n, p, s, t) and Kruskal-Wallis test (q). *P < 0.05, **P < 0.01, ***P < 0.001.

**Supplementary Figure 2.**
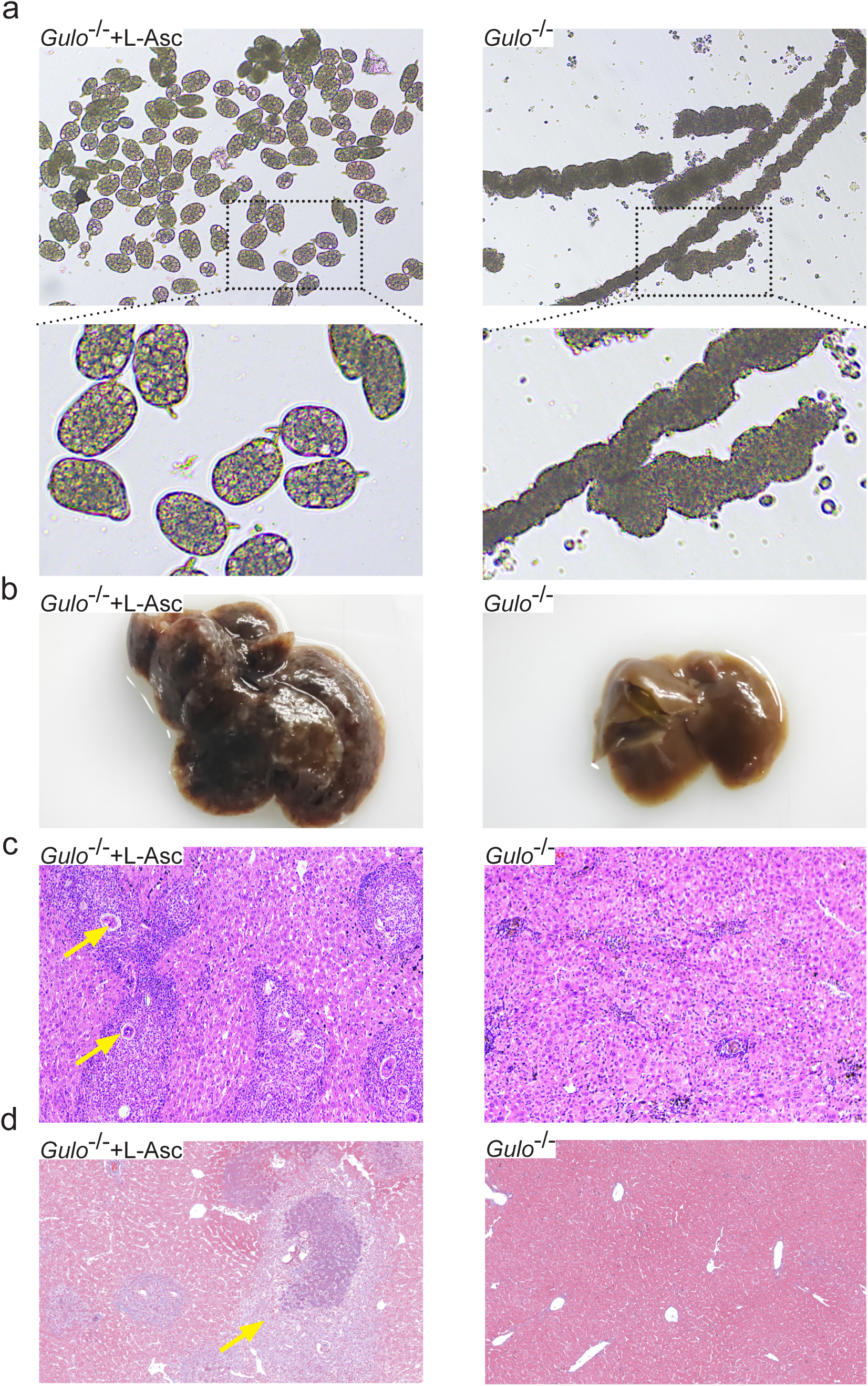
Effects of ascorbate deficiency on egg production and pathogenesis of *S. japonicum* infection. **a.** Eggs produced by *S. japonicum* worms isolated from infected *Gulo^-/-^* mice that were supplemented or not supplemented with ascorbate. Eggs were analyzed 1 day after worm isolation from mice and culture in BM169 media without ascorbate. **b.** Liver enlargement and granulomas in ascorbate-supplemented but not unsupplemented *S. japonicum-*infected *Gulo^-/-^* mice **c.** H & E showing granulomas forming around an egg (yellow arrow) in ascorbate-supplemented but not unsupplemented *S. japonicum-*infected *Gulo^-/-^* mice, **d.** Masson’s Trichrome staining showing liver fibrosis (yellow arrow) in ascorbate-supplemented but not unsupplemented *S. japonicum-*infected *Gulo^-/-^* mice.

**Supplementary Figure 3.**
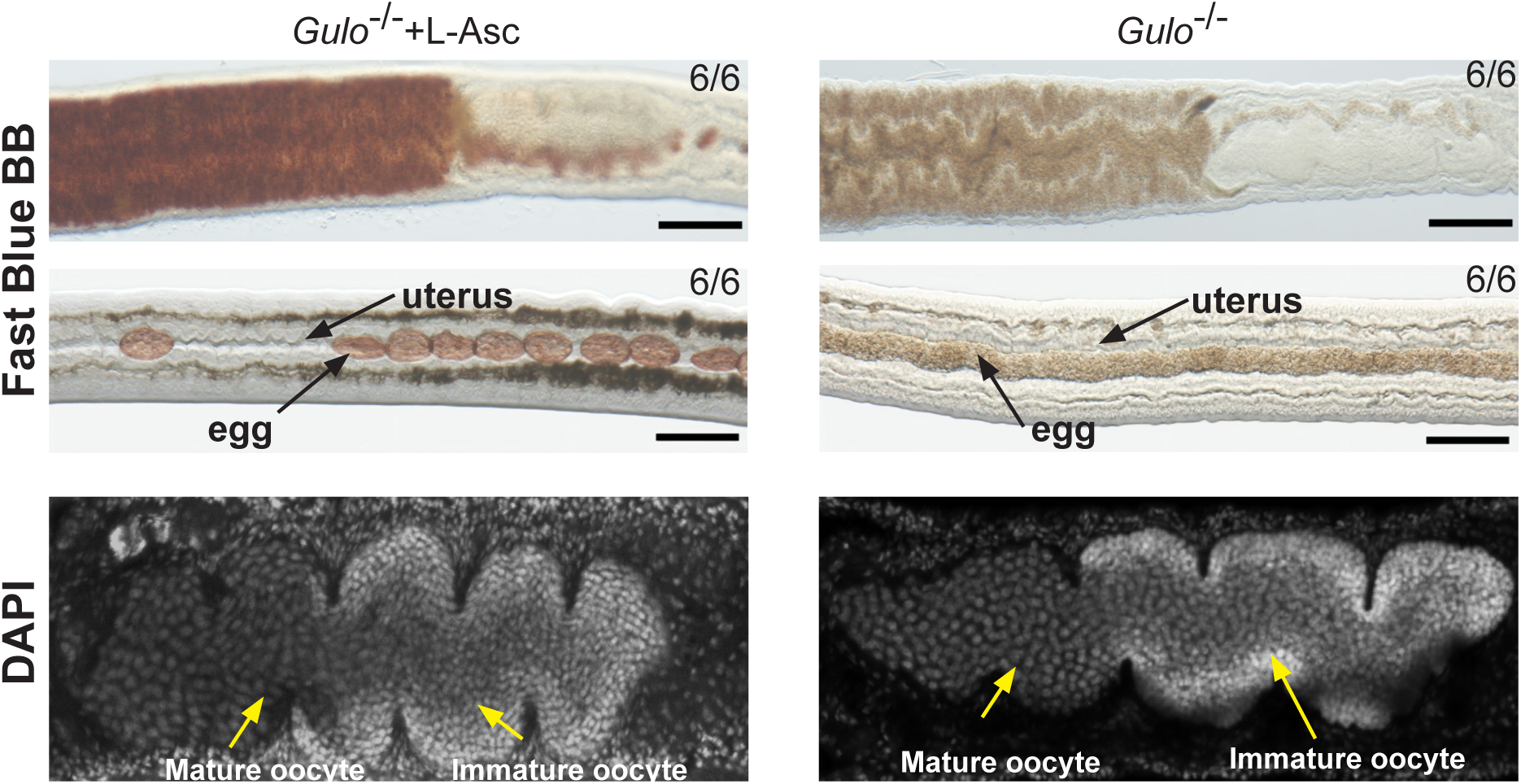
Reproductive development of female *S. japonicum* isolated from ascorbate supplemented or unsupplemented *Gulo*^-/-^ mice. Fast Blue BB staining showing the vitellaria (top) and eggs (middle) and DAPI staining showing the ovaries (bottom) of adult female parasites from ascorbate-depleted or supplemented *Gulo*^-/-^ mice following paired infections.

**Supplementary Figure 4.**
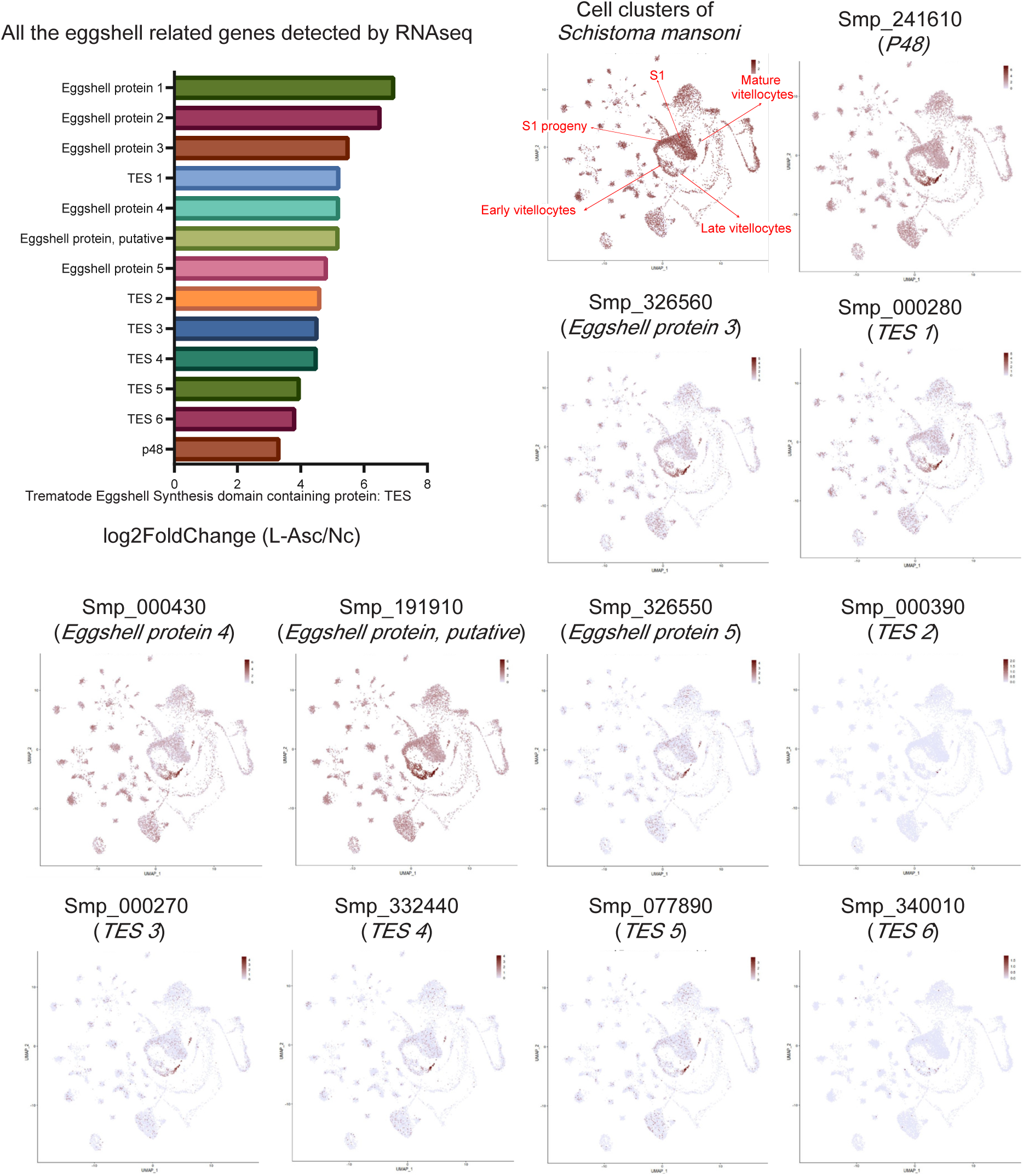
Expression of eggshell synthesis genes is stimulated by ascorbate treatment. Examples of genes involved in vitellocyte development and eggshell synthesis whose expression is stimulated by ascorbate. Data from RNA-seq experiment shown in Fig. 4.

**Supplementary Figure 5.**
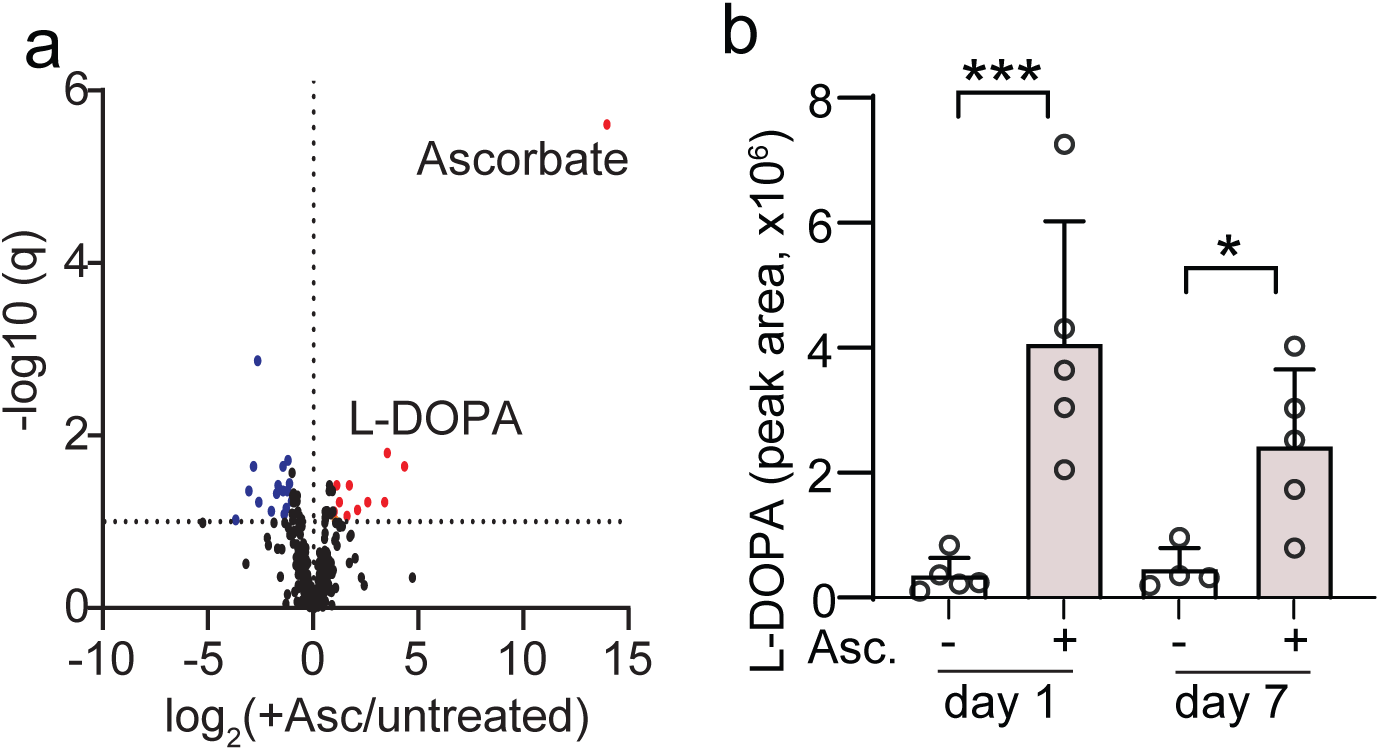
Metabolomics analysis of ascorbate-treated female *S. mansoni*. **a.** Metabolomics analysis of paired female worms treated with 200 μM L-Ascorbate for 24 hours as compared to untreated controls. Each dot represents a metabolite. Statistical significance was assessed with multiple t-tests with false discovery rate correction. **b.** Elevated L-DOPA in ascorbate-treated female worms. n = 4-5 samples/treatment, with 5 worms/sample.

**Supplementary Figure 6.**
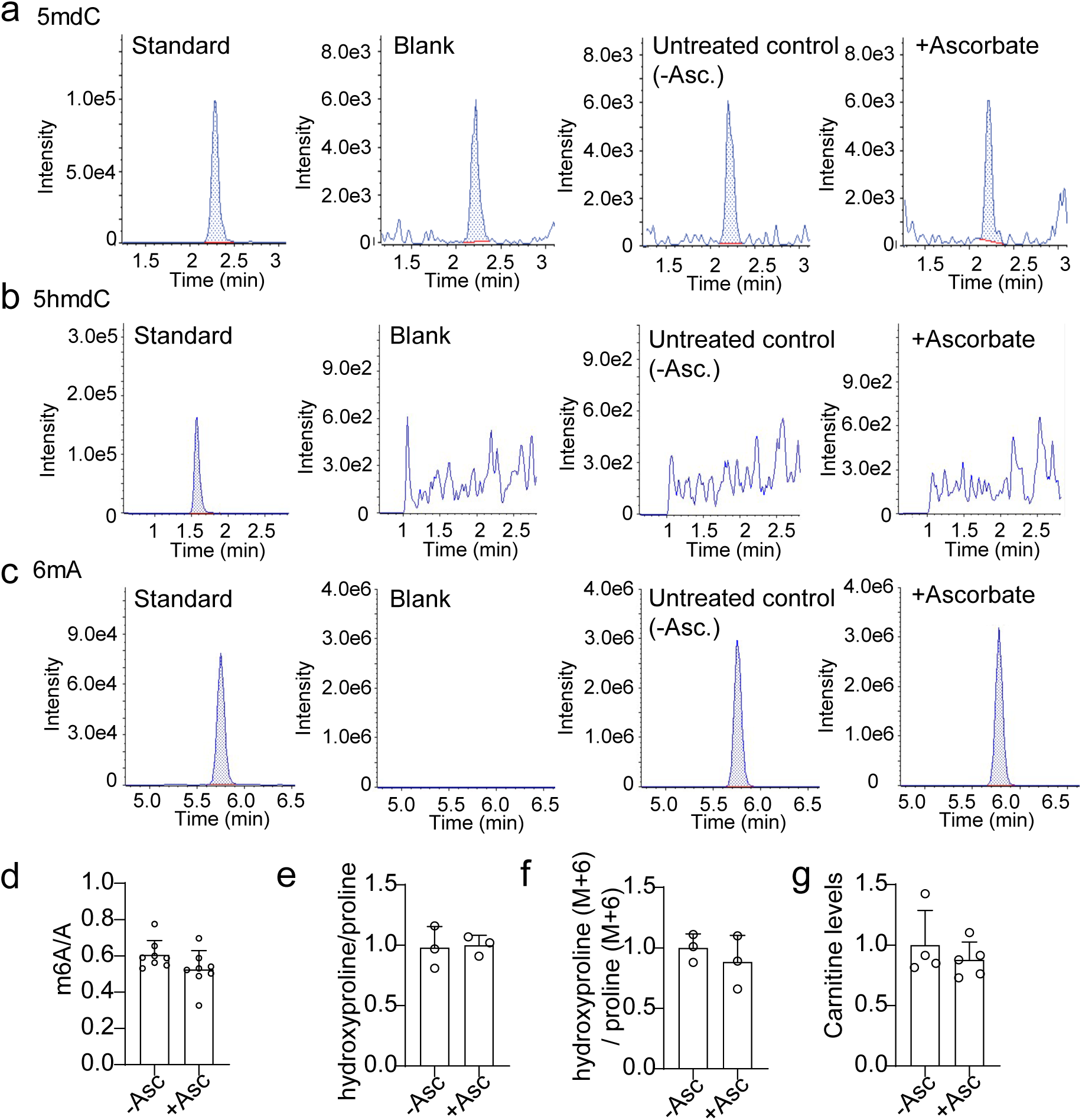
Levels of reaction products of select dioxygenase enzymes in ascorbate-treated female *S. mansoni*. **a-c.** LC-MS/MS chromatogram (MRM) for 5mdC (5-methyl-2’-deoxycytidine), 5hmdC (5-hydroxymethyl-2’-deoxycytidine), or 6mA (N6-Methyladenosine) detected from chemical standards, or enzymatic digestion blanks, or female worms paired and incubated for 1 week in BC169 media with (+Ascorbate) or without (-Ascorbate) 200 μM ascorbate. **d.** Levels of m6A in mRNA. m6A and A from hydrolyzed mRNA were quantified in ascorbate-treated (+Asc) or untreated (-Asc) female worms by LC-MS/MS with m6A-^13^C1 and A-^13^C5 labeled isotopes as internal standards. n = 8 biological replicates with 5 worms/replicate. **e.** Measurement of proline hydroxylation. The ratio of hydroxyproline and proline peak area in hydrolyzed proteins was measured by LC/MS in paired female worms after 7 days of incubation with or without 200 μM ascorbate. n = 3 biological replicates with 5 worms/replicate. **f.** Measurement of newly hydroxylated proline on proteins. Paired worms were incubated for 4 days with ^13^C5,^15^N-L-Proline (M+6), with or without 200 μM ascorbate, and levels of ^13^C5,^15^N-hydroxyroline (M+6) and ^13^C5,^15^N-L-Proline (M+6) in hydrolyzed proteins from female worms were measured by LC/MS. n = 3 biological replicates. **g.** Measurement of carnitine levels. Carnitine levels were measured by LC/MS in female worms after 7 days of treatment with or without 200 μM ascorbate. n = 4-5 biological replicates. All graphs show means ± sd. Statistical significance was assessed unpaired t-test (d-g).

**Supplementary Figure 7.**
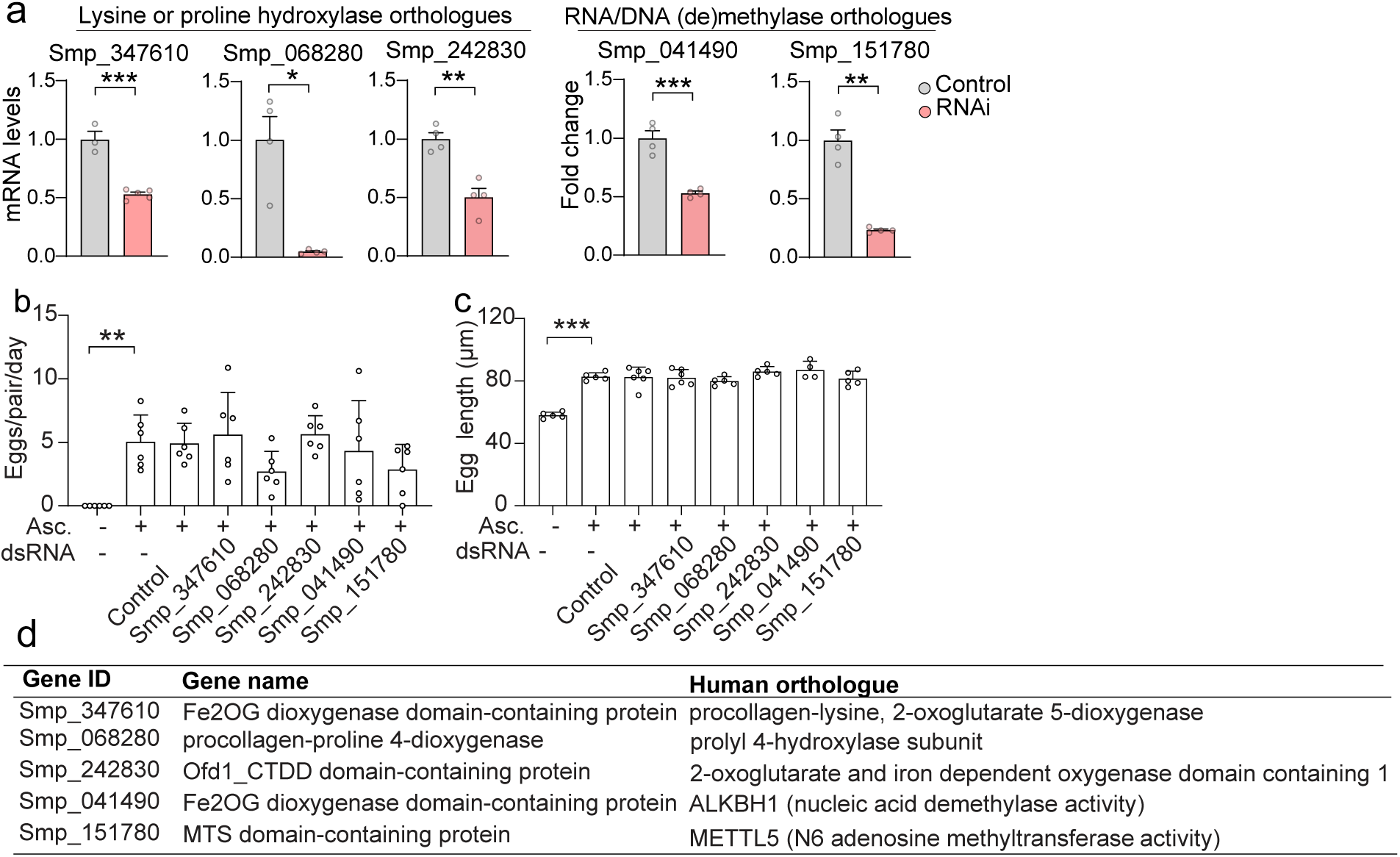
Inhibition of candidate nucleic acid and protein modification enzymes does not affect *S. mansoni* egg production. **a.** mRNA levels of genes involved in proline/lysine hydroxylation or nucleic acid methylation after 20 days of incubation with corresponding dsRNA. n = 3-5 biological replicates with 5 worms/replicate. **b, c.** Effects of RNAi against indicated genes on egg length and number. The number of normal eggs per day per pair of worms between day 16 and day 18 of dsRNA with 200 μM ascorbate treatment was counted. Incubation without ascorbate is shown as a negative control. n = 5-6 biological replicates. **d.** Annotation of *S. mansoni* genes inhibited by RNAi. All graphs show means ± sd. Statistical significance was assessed with unpaired t-test (a: Smp_347610, Smp_242830, Smp_041490), Welch’s test (a: Smp_068280, Smp_151780), Kruskal-Wallis test (b) and one-way ANOVA (c). *P < 0.05, **P < 0.01, ***P < 0.001.

**Supplementary Figure 8.**
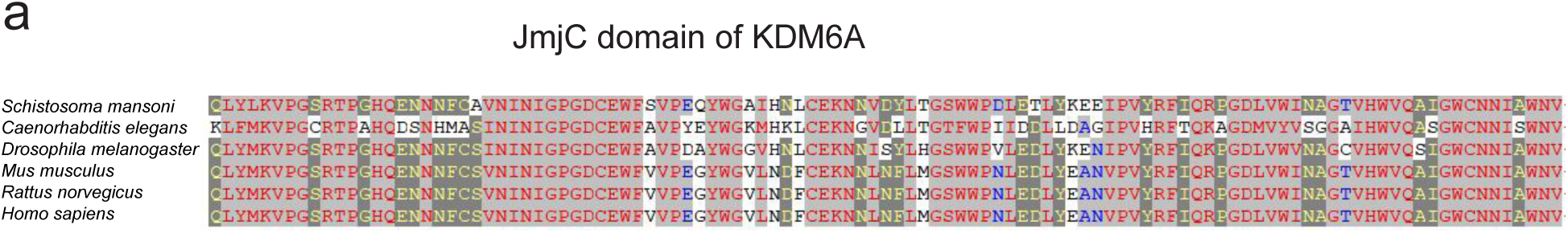
Conservation of *S. mansoni* KDM6 JmjC domain. Alignment of the Kdm6 JmjC domain in *S. mansoni* and model organisms.

**Supplementary Figure 9.**
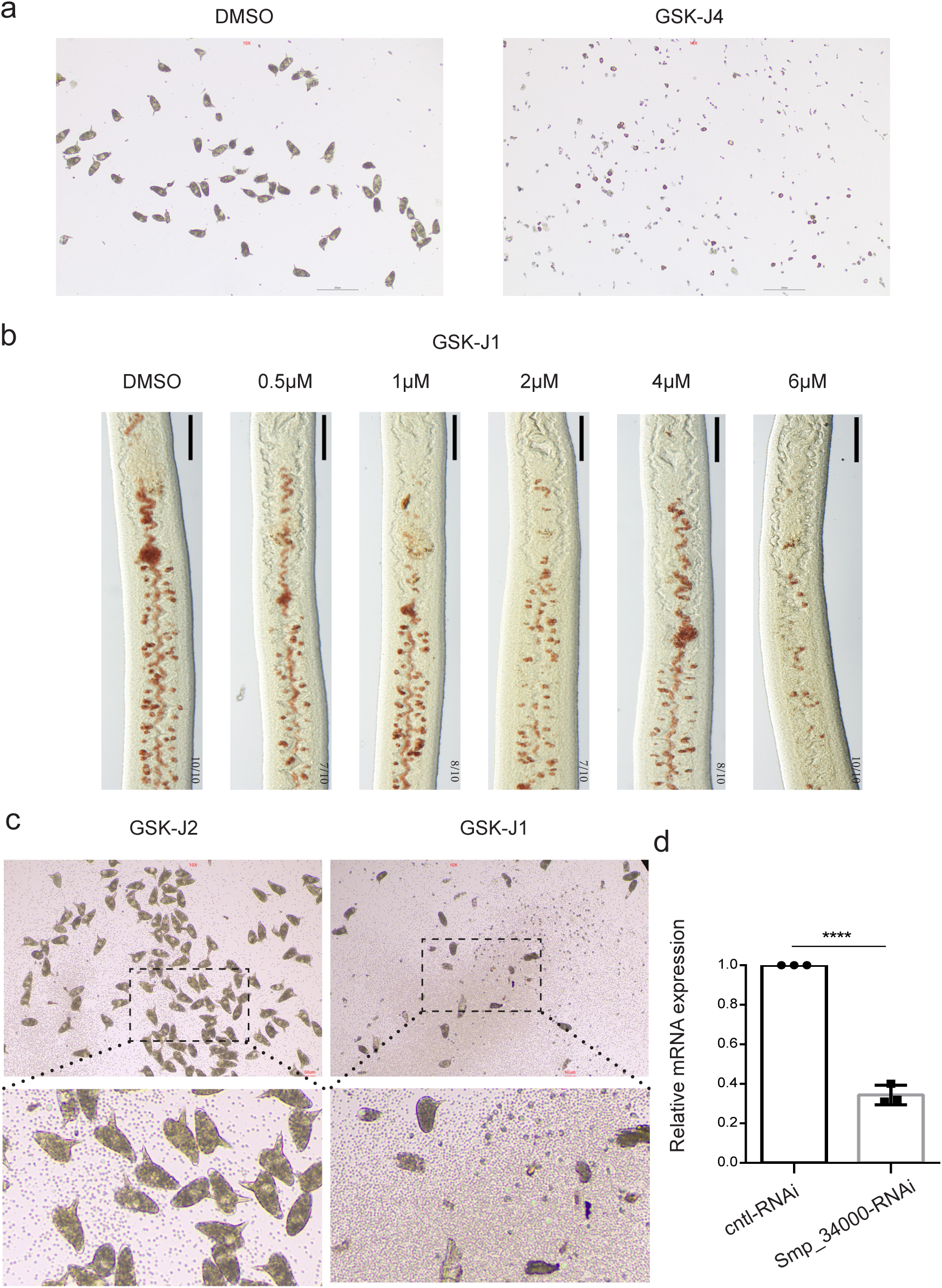
Effects of KDM inhibition on egg laying and vitellocyte development. **a.** Eggs laid by paired adult females treated with GSK-J4. **b.** Titration of GSK-J1 and effects on vitellaria development of virgin females treated with BATT. Vitellaria were stained by Fast blue BB staining. **c.** Eggs laid by paired adult females treated with GSK-J1 or GSK-J2. **d.** Reduction in *Smp_034000* mRNA levels after RNAi.

**Supplementary Table 3.**
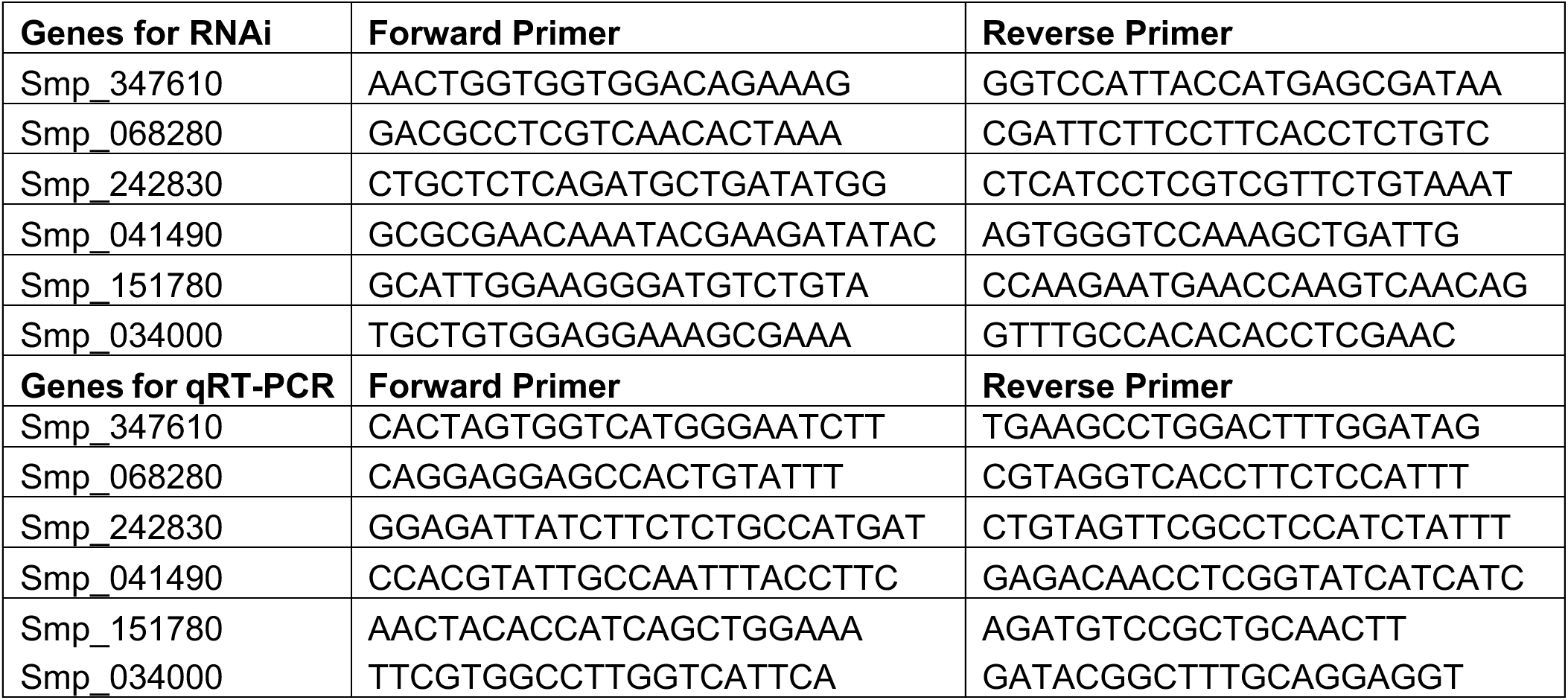
Primers used in this study.

## References

1 Wheeler, G., Ishikawa, T., Pornsaksit, V. & Smirnoff, N. Evolution of alternative biosynthetic pathways for vitamin C following plastid acquisition in photosynthetic eukaryotes. Elife 4 (2015). 10.7554/eLife.06369

2 Albalat, R. & Canestro, C. Evolution by gene loss. Nat Rev Genet 17, 379–391 (2016). 10.1038/nrg.2016.39

3 Schleicher, R. L., Carroll, M. D., Ford, E. S. & Lacher, D. A. Serum vitamin C and the prevalence of vitamin C deficiency in the United States: 2003-2004 National Health and Nutrition Examination Survey (NHANES). Am J Clin Nutr 90, 1252–1263 (2009). 10.3945/ajcn.2008.27016

4 Khaw, K. T. et al. Relation between plasma ascorbic acid and mortality in men and women in EPIC-Norfolk prospective study: a prospective population study. European Prospective Investigation into Cancer and Nutrition. Lancet 357, 657–663 (2001).

5 Loria, C. M., Klag, M. J., Caulfield, L. E. & Whelton, P. K. Vitamin C status and mortality in US adults. Am J Clin Nutr 72, 139–145 (2000).

6 Frei, B., England, L. & Ames, B. N. Ascorbate is an outstanding antioxidant in human blood plasma. Proc Natl Acad Sci U S A 86, 6377–6381 (1989). 10.1073/pnas.86.16.6377

7 Linster, C. L. & Van Schaftingen, E. Vitamin C. Biosynthesis, recycling and degradation in mammals. FEBS J 274, 1–22 (2007). 10.1111/j.1742-4658.2006.05607.x

8 Agathocleous, M. et al. Ascorbate regulates haematopoietic stem cell function and leukaemogenesis. Nature 549, 476–481 (2017). 10.1038/nature23876

9 DiTroia, S. P. et al. Maternal vitamin C regulates reprogramming of DNA methylation and germline development. Nature 573, 271–275 (2019). 10.1038/s41586-019-1536-1

10 Thaler, R. et al. Vitamin C epigenetically controls osteogenesis and bone mineralization. Nat Commun 13, 5883 (2022). 10.1038/s41467-022-32915-8

11 Bornstein, S. R. et al. Impaired adrenal catecholamine system function in mice with deficiency of the ascorbic acid transporter (SVCT2). FASEB J 17, 1928–1930 (2003). 10.1096/fj.02-1167fje

12 Maeda, N. et al. Aortic wall damage in mice unable to synthesize ascorbic acid. Proc Natl Acad Sci U S A 97, 841–846 (2000). 10.1073/pnas.97.2.841

13 Agathocleous, M. The physiological functions of ascorbate in the development of cancer. Dis Model Mech 18 (2025). 10.1242/dmm.052201

14 Carpenter, K. J. The History of Scurvy and Vitamin C. (Cambridge University Press, 1988).

15 Snoddy, A. M. E. et al. Macroscopic features of scurvy in human skeletal remains: A literature synthesis and diagnostic guide. Am J Phys Anthropol 167, 876–895 (2018). 10.1002/ajpa.23699

16 McManus, D. P. et al. Schistosomiasis. Nat Rev Dis Primers 4, 13 (2018). 10.1038/s41572-018-0013-8

17 King, C. H. Parasites and poverty: the case of schistosomiasis. Acta Trop 113, 95–104 (2010). 10.1016/j.actatropica.2009.11.012

18 Colley, D. G., Bustinduy, A. L., Secor, W. E. & King, C. H. Human schistosomiasis. Lancet 383, 2253–2264 (2014). 10.1016/S0140-6736(13)61949-2

19 Pearce, E. J. & MacDonald, A. S. The immunobiology of schistosomiasis. Nat Rev Immunol 2, 499–511 (2002). 10.1038/nri843

20 Takaki, K. K., Rinaldi, G., Berriman, M., Pagan, A. J. & Ramakrishnan, L. Schistosoma mansoni Eggs Modulate the Timing of Granuloma Formation to Promote Transmission. Cell Host Microbe 29, 58–67 e55 (2021). 10.1016/j.chom.2020.10.002

21 Abdel Aziz, N., Musaigwa, F., Mosala, P., Berkiks, I. & Brombacher, F. Type 2 immunity: a two-edged sword in schistosomiasis immunopathology. Trends Immunol 43, 657–673 (2022). 10.1016/j.it.2022.06.005

22 Le Clec’h, W. et al. Genetic analysis of praziquantel response in schistosome parasites implicates a transient receptor potential channel. Sci Transl Med 13, eabj9114 (2021). 10.1126/scitranslmed.abj9114

23 Chen, R. et al. A male-derived nonribosomal peptide pheromone controls female schistosome development. Cell 185, 1506–1520 e1517 (2022). 10.1016/j.cell.2022.03.017

24 Wang, J. & Collins, J. J., 3rd. Identification of new markers for the Schistosoma mansoni vitelline lineage. Int J Parasitol 46, 405–410 (2016). 10.1016/j.ijpara.2016.03.004

25 Krakower, C., Hoffman, W. A. & Axtmayer, J. H. Defective Granular Eggshell Formation by Schistosoma Mansoni in Experimentally Infected Guinea Pigs on a Vitamin C Deficient Di. The Journal of Infectious Diseases 74, 178–183 (1944).

26 Wang, J., Chen, R. & Collins, J. J., 3rd. Systematically improved in vitro culture conditions reveal new insights into the reproductive biology of the human parasite Schistosoma mansoni. PLoS Biol 17, e3000254 (2019). 10.1371/journal.pbio.3000254

27 You, Y. et al. An improved medium for in vitro studies of female reproduction and oviposition in Schistosoma japonicum. Parasit Vectors 17, 116 (2024). 10.1186/s13071-024-06191-y

28 Wijshake, T. et al. Schistosome Infection Impacts Hematopoiesis. J Immunol 212, 607–616 (2024). 10.4049/jimmunol.2300195

29 Oguro, H., Ding, L. & Morrison, S. J. SLAM family markers resolve functionally distinct subpopulations of hematopoietic stem cells and multipotent progenitors. Cell Stem Cell 13, 102–116 (2013). 10.1016/j.stem.2013.05.014

30 Pietras, E. M. et al. Functionally Distinct Subsets of Lineage-Biased Multipotent Progenitors Control Blood Production in Normal and Regenerative Conditions. Cell Stem Cell 17, 35–46 (2015). 10.1016/j.stem.2015.05.003

31 Chuah, C., Jones, M. K., Burke, M. L., McManus, D. P. & Gobert, G. N. Cellular and chemokine-mediated regulation in schistosome-induced hepatic pathology. Trends Parasitol 30, 141–150 (2014). 10.1016/j.pt.2013.12.009

32 Crandon, J. H., Lund, C. C. & Dill, D. B. Experimental Human Scurvy. N Engl J Med 223, 353–369 (1940).

33 Krebs, H. A. The Sheffield Experiment on the Vitamin C Requirement of Human Adults. Proceedings of the Nutrition Society 12, 237–246 (1953). 10.1079/PNS19530054

34 Wendt, G. et al. A single-cell RNA-seq atlas of Schistosoma mansoni identifies a key regulator of blood feeding. Science 369, 1644–1649 (2020). 10.1126/science.abb7709

35 Flashman, E., Davies, S. L., Yeoh, K. K. & Schofield, C. J. Investigating the dependence of the hypoxia-inducible factor hydroxylases (factor inhibiting HIF and prolyl hydroxylase domain 2) on ascorbate and other reducing agents. Biochem J 427, 135–142 (2010). 10.1042/BJ20091609

36 Minor, E. A., Court, B. L., Young, J. I. & Wang, G. Ascorbate induces ten-eleven translocation (Tet) methylcytosine dioxygenase-mediated generation of 5-hydroxymethylcytosine. J Biol Chem 288, 13669–13674 (2013). 10.1074/jbc.C113.464800

37 Blaschke, K. et al. Vitamin C induces Tet-dependent DNA demethylation and a blastocyst-like state in ES cells. Nature 500, 222–226 (2013). 10.1038/nature12362

38 Chen, J. et al. Vitamin C modulates TET1 function during somatic cell reprogramming. Nat Genet 45, 1504-1509 (2013). 10.1038/ng.2807

39 Myllyla, R., Majamaa, K., Gunzler, V., Hanauske-Abel, H. M. & Kivirikko, K. I. Ascorbate is consumed stoichiometrically in the uncoupled reactions catalyzed by prolyl 4-hydroxylase and lysyl hydroxylase. J Biol Chem 259, 5403–5405 (1984).

40 Young, J. I., Zuchner, S. & Wang, G. Regulation of the Epigenome by Vitamin C. Annu Rev Nutr 35, 545–564 (2015). 10.1146/annurev-nutr-071714-034228

41 Whatley, K. C. L. et al. The repositioning of epigenetic probes/inhibitors identifies new anti-schistosomal lead compounds and chemotherapeutic targets. PLoS Negl Trop Dis 13, e0007693 (2019). 10.1371/journal.pntd.0007693

42 Lobo-Silva, J. et al. The antischistosomal potential of GSK-J4, an H3K27 demethylase inhibitor: insights from molecular modeling, transcriptomics and in vitro assays. Parasit Vectors 13, 140 (2020). 10.1186/s13071-020-4000-z

43 Kruidenier, L. et al. A selective jumonji H3K27 demethylase inhibitor modulates the proinflammatory macrophage response. Nature 488, 404–408 (2012). 10.1038/nature11262

44 Parker, S. J. et al. Spontaneous hydrolysis and spurious metabolic properties of alpha-ketoglutarate esters. Nat Commun 12, 4905 (2021). 10.1038/s41467-021-25228-9

45 Heinemann, B. et al. Inhibition of demethylases by GSK-J1/J4. Nature 514, E1–2 (2014). 10.1038/nature13688

46 Long, C. L. et al. Ascorbic acid dynamics in the seriously ill and injured. J Surg Res 109, 144–148 (2003).

47 Marcus, S. L. et al. Hypovitaminosis C in patients treated with high-dose interleukin 2 and lymphokine-activated killer cells. Am J Clin Nutr 54, 1292S–1297S (1991). 10.1093/ajcn/54.6.1292s

48 Du, W. D. et al. Therapeutic efficacy of high-dose vitamin C on acute pancreatitis and its potential mechanisms. World J Gastroenterol 9, 2565–2569 (2003). 10.3748/wjg.v9.i11.2565

49 Chevion, S., Or, R. & Berry, E. M. The antioxidant status of patients subjected to total body irradiation. Biochem Mol Biol Int 47, 1019–1027 (1999).

50 Fowler, A. A., 3rd et al. Phase I safety trial of intravenous ascorbic acid in patients with severe sepsis. J Transl Med 12, 32 (2014). 10.1186/1479-5876-12-32

51 Galley, H. F., Davies, M. J. & Webster, N. R. Ascorbyl radical formation in patients with sepsis: effect of ascorbate loading. Free Radic Biol Med 20, 139–143 (1996).

52 Crandon, J. H. et al. Ascorbic acid economy in surgical patients as indicated by blood ascorbic acid levels. N Engl J Med 258, 105–113 (1958). 10.1056/NEJM195801162580301

53 Rowe, S. & Carr, A. C. Global Vitamin C Status and Prevalence of Deficiency: A Cause for Concern? Nutrients 12 (2020). 10.3390/nu12072008

54 Kariuki, S. N. & Williams, T. N. Human genetics and malaria resistance. Hum Genet 139, 801–811 (2020). 10.1007/s00439-020-02142-6

55 Diaz, A. V., Walker, M. & Webster, J. P. Reaching the World Health Organization elimination targets for schistosomiasis: the importance of a One Health perspective. Philos Trans R Soc Lond B Biol Sci 378, 20220274 (2023). 10.1098/rstb.2022.0274

56 Shehata, M. A., Chama, M. F. & Funjika, E. Prevalence and intensity of Schistosoma haematobium infection among schoolchildren in central Zambia before and after mass treatment with a single dose of praziquantel. Trop Parasitol 8, 12–17 (2018). 10.4103/tp.TP_32_17

57 Flammer, P. G. et al. Epidemiological insights from a large-scale investigation of intestinal helminths in Medieval Europe. PLoS Negl Trop Dis 14, e0008600 (2020). 10.1371/journal.pntd.0008600

58 Standley, C. J., Mugisha, L., Dobson, A. P. & Stothard, J. R. Zoonotic schistosomiasis in non-human primates: past, present and future activities at the human-wildlife interface in Africa. J Helminthol 86, 131–140 (2012). 10.1017/S0022149X12000028

59 Helliwell, K. E., Wheeler, G. L. & Smith, A. G. Widespread decay of vitamin-related pathways: coincidence or consequence? Trends Genet 29, 469–478 (2013). 10.1016/j.tig.2013.03.003

60 Collins, J. J., 3rd et al. Adult somatic stem cells in the human parasite Schistosoma mansoni. Nature 494, 476–479 (2013). 10.1038/nature11924

61 Wang, J. et al. Large-scale RNAi screening uncovers therapeutic targets in the parasite Schistosoma mansoni. Science 369, 1649–1653 (2020). 10.1126/science.abb7699

62 Jun, S. et al. The requirement for pyruvate dehydrogenase in leukemogenesis depends on cell lineage. Cell Metab 33, 1777–1792 e1778 (2021). 10.1016/j.cmet.2021.07.016

